# CellScope: High-Performance Cell Atlas Workflow with Tree-Structured Representation

**DOI:** 10.1101/2025.02.15.638400

**Authors:** Bingjie Li, Runyu Lin, Tianhao Ni, Guanao Yan, Mannix Burns, Jingyi Jessica Li, Zhigang Yao

## Abstract

Single-cell sequencing enables comprehensive profiling of individual cells, revealing cellular heterogeneity and function with unprecedented resolution. However, current analysis frameworks lack the ability to simultaneously explore and visualize cellular hierarchies at multiple biological levels. To address these limitations, we present CellScope, an innovative framework for constructing high-resolution cell atlases at multiple clustering levels. CellScope employs a two-step manifold fitting process for gene selection and noise reduction, followed by agglomerative clustering, and uniquely integrates UMAP visualization with hierarchical clustering to intuitively represent cellular relationships simultaneously at multiple levels—such as cell lineage, cell type, and cell subtype levels. Compared to established pipelines such as Seurat and Scanpy, CellScope comprehensively improves clustering performance, visualization clarity, computational efficiency, and algorithm interpretability, while reducing dependence on hyperparameters across a multitude of single-cell datasets. Most importantly, it can reveal new biological insights that other contemporary methods are unable to detect, thereby deepening our understanding of cellular heterogeneity and function, and potentially informing disease research.

## Introduction

The advent of single-cell sequencing has fundamentally changed our understanding of biology by providing an unprecedented look into the heterogeneity of biological systems at the individual cell level. Over the past decade, the increasing accessibility of single-cell technologies has led to a rise in the generation of large comprehensive single-cell datasets-collectively known as cell atlases. By providing comprehensive high-resolution maps that identify, characterize, and spatially locate every cell type within an organism or tissue, cell atlases offer invaluable insights into cellular heterogeneity, interactions, and functions [1]. These detailed maps have the potential to revolutionize our understanding of normal development, aging, and disease pathogenesis, paving the way for new diagnostic, prognostic, and therapeutic strategies [2, 3]. Consequently, many specialized atlases have emerged, including those focusing on neurodegenerative diseases-mapping cellular changes in Alzheimer’s and Parkinson’s disease tissues [4, 5, 6]. Others have focused on developmental biology, creating time-resolved atlases that track cellular differentiation during organ formation [1, 7, 8]. Cancer-specific atlases have also gained prominence, helping to delineate tumor micro-environments and identify new therapeutic targets [9, 10]. With this increasing availability of cell atlases, the ability to extract biologically meaningful information from these datasets is paramount to the progression of our knowledge of system-specific cellular dynamics and disease mechanisms.

Despite remarkable progress in the single-cell field, existing computational methodologies face several limitations that hinder their ability to fully capture the complexity of single-cell data. Commonly used pipelines, such as Seurat [11], Scanpy [12], and SnapATAC [13], rely on conventional unsupervised learning techniques that may not adequately handle the high-dimensionality, sparsity, and noise inherent in single-cell datasets. For instance, the highly variable genes (HVG) selection method employed by Seurat can be sensitive to technical noise and may overlook rare cell types with subtle but biologically relevant variations [14]. Similarly, the Louvain clustering algorithm, widely used in pipelines, struggles to identify small cell populations and lacks a hierarchical structure that captures the nested relationships and developmental trajectories between cell types [15]. Moreover, popular visualization techniques like t-SNE and UMAP have limitations in representing the global structure and hierarchical organization of cells, often emphasizing local similarities at the expense of preserving the overall topology [16].

Recent advances in single-cell genomics have led to growing recognition of the low-dimensional nature of single-cell data [17, 18]. This characteristic can be understood from two perspectives. Firstly, despite the vast number of genes measured in single-cell experiments, only a small subset is typically informative for distinguishing cell types. The majority of genes are housekeeping genes, which maintain basic cellular functions and exhibit relatively constant expression across cell types. Studies such as [14] have demonstrated that focusing on fewer, highly informative genes can lead to improved visualization and analysis outcomes [19]. Secondly, due to the interconnected nature of genes, single-cell data tends to occupy a low-dimensional manifold within the high-dimensional gene expression space [20]. As such, many state-of-the-art single-cell clustering frameworks have begun incorporating this concept of manifolds [21, 22]. In particular, the process of manifold fitting [23, 24], which preserves data structure while still offering high interpretability and theoretical backing, has emerged as a superior dimensionality reduction technique. Recent advancements in [24] have led to the development of scAMF [25]—the first framework to apply manifold fitting to single-cell analysis.

Here, we introduce “CellScope”, an innovative method for constructing multi-level, high-resolution cellular atlases. By leveraging manifold fitting and neighborhood graph-based aggregative clustering, CellScope addresses key challenges in single-cell analysis. We show that CellScope excels in three critical areas: (1) precise delineation of similar cell types and detection of rare populations, (2) dynamic characterization of cellular landscapes at multiple cell type and subtype resolutions, and (3) multi-level functional analysis of genes. We conducted extensive validation across 36 datasets covering various species, organs, and sequencing modalities such as scRNA-seq and scATAC-seq, demonstrating CellScope’s exceptional clustering performance. Its capabilities are further illustrated by the identification and functional characterization of novel oligodendrocyte subpopulations, as well as simultaneous detection of cell types and health status in peripheral blood cells of COVID-19 patients, neither of which were uncovered by existing methods, such as Seurat and Scanpy. Importantly, CellScope achieves these results with exceptional speed, adaptive parameter-free operation, and consistent performance, while maintaining high interpretability. This innovative approach opens new avenues for comprehending cellular diversity and function in complex biological systems.

## Results

### Overview of CellScope workflow

CellScope uses manifold fitting to model single-cell data, tackling the hurdles of intrinsic complexity and noise to derive crucial biological insights. CellScope assumes that the true biological structure of single-cell data lies on a low-dimensional manifold [26]. This manifold represents the intrinsic, lower-dimensional structure of gene expression that captures the genuine relationships between cells, including cellular states and subtypes. However, the observed single-cell data does not directly reflect this manifold due to two types of noise. The first type of noise refers to the expression of housekeeping genes, which are crucial for basic cellular functions but, due to their ubiquitous expression, do not contribute to distinguishing cell populations. We thus use “*noise space*” to denote the space of housekeeping genes and “*signal space*” to represent the remaining gene expression profiles that reflect cell type differences. The second type of noise lies in this signal space and represents technical noise due to mRNA loss, inefficient molecule capturing, and sequencing errors [27]. This stochastic noise may distort the true expression patterns of the marker genes that are key to distinguishing between cell types and states. Together, these two types of noise combine with the underlying biologically meaningful manifold to constitute the observable single-cell gene expression matrix (**Figure 1a**).

**Figure 1.**
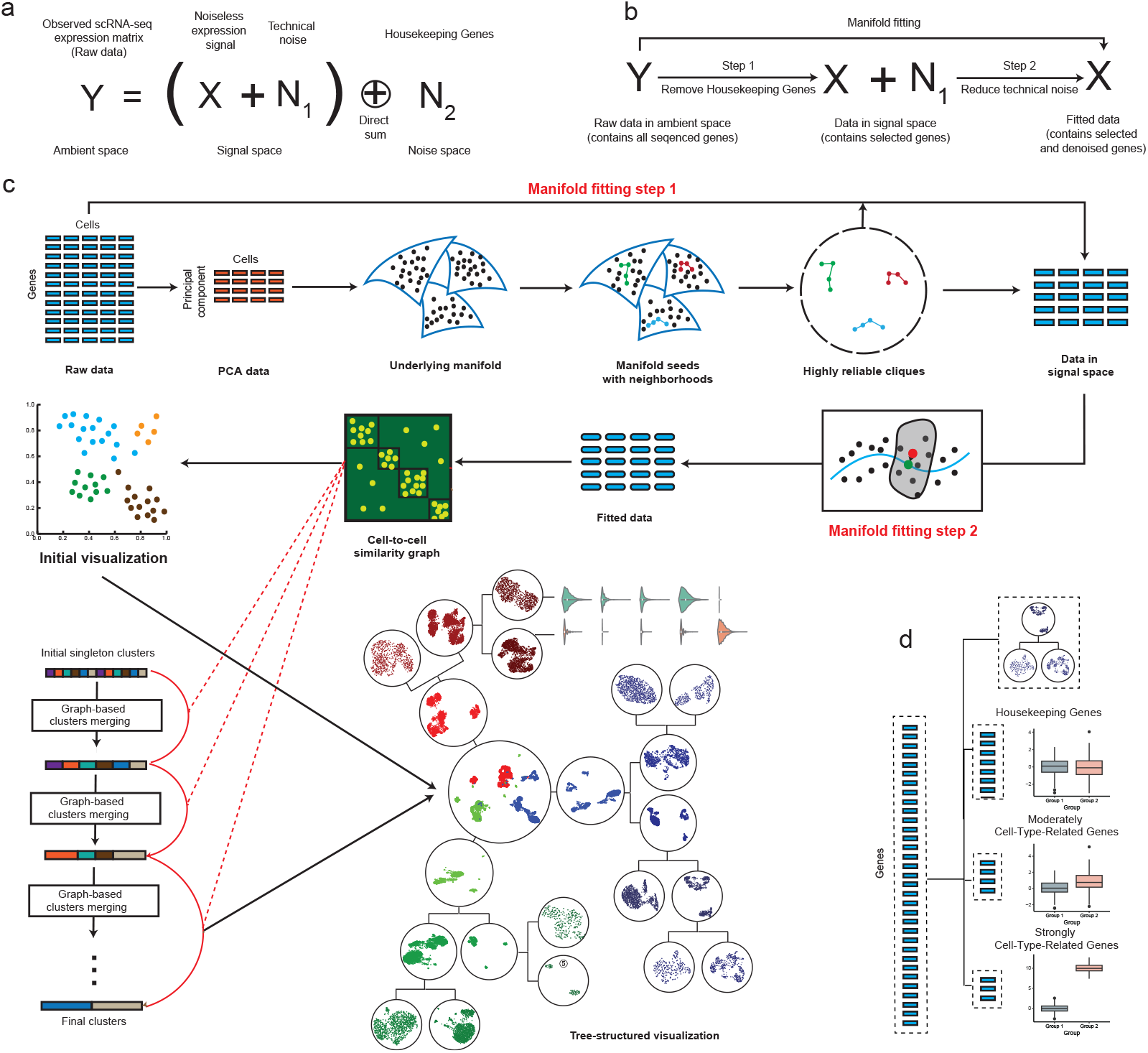
**a**. Mathematical modeling of noise in single-cell data. **b**. The two steps of manifold fitting and their purpose. **c**. Overview of CellScope workflow. CellScope enhances cellular data analysis through a two-step manifold fitting process. First, it identifies “manifold seeds” and “highly reliable cliques” in the PCA-reduced space to effectively distinguish signal from noise, thereby removing housekeeping genes. Next, it reduces technical noise by projecting low-density cells onto high-density regions. Subsequently, CellScope constructs a neighborhood similarity graph and performs agglomerative clustering, iteratively merging similar clusters for precise hierarchical classification. Finally, the method generates two key visualizations: the UMAP of the manifold-fitted data and a tree-structured visualization combining UMAP with hierarchical clustering. **d**. Each cluster divisions in the tree-structured visualization produce three unique types of genes: Housekeeping Genes with minimal variance between classes, Moderately Cell-Type-Related Genes with partially significant differences, and Strongly Cell-Type-Related Genes with large diversity.

To recover the essential low-dimensional manifold and improve the quality of downstream analyses, CellScope employs a two-step manifold fitting process **(Figure 1b)**. The first step in our manifold fitting approach aims to mitigate the noise introduced by the inconsistent expression of housekeeping genes, which are irrelevant to cell classification, while preserving critical genes for further analysis **(Methods A)**. We base this process on a widely accepted assumption in manifold learning and translate it to a biological context: low-dimensional representations of individual cells that belong to different cell-types lie on distinct submanifolds [28]. These cell-type submanifolds are characterized by a high density of cells and are separated from one another by regions of low cell density. [29].

Leveraging this principle, CellScope selects multiple sets of distant high-density cells, termed “manifold seeds”, along with their neighboring cells, designated as “highly reliable cliques” (**Figure 1c**). These cliques originate from multiple separate cell-types and help distinguish between noise and signal spaces. Features in the signal space exhibit low variance within the same clique but high variance between different cliques, while features in the noise space lack this property. By exploiting this distinction, we filter out most noise while preserving key genes for determining cell identity.

The second step ensures proper stratification of different cell-types by assigning cells residing in low-density regions, which may represent transitional cell states or have higher levels of technical noise, to the nearest cell-type submanifold **(Methods B)**. This denoising step refines the representation of each cell-type, emphasizing genuine biological signals over technical artifacts, and better reflects the underlying cellular heterogeneity.

After manifold fitting, CellScope constructs a cell-to-cell neighborhood similarity graph, where cells with more similar gene expression profiles are assigned higher similarity. Based on this graph, CellScope then performs agglomerative clustering **(Methods C)**. Starting with each cell as a cluster, the algorithm iteratively merges the most similar clusters until no two clusters exhibit significant similarity, yielding precise and biologically meaningful classifications.

A novel aspect of CellScope is that CellScope generates an informative tree-structured diagram **(Methods D)** that integrates UMAP [30] and hierarchical clustering. In addition to an initial UMAP visualization of the manifold-fitted data that provides an intuitive representation of complex cellular relationships, it provides the tree-structured visualization that depicts the hierarchical relationships between cell types, illustrating how different populations emerge, branch, and specialize. By annotating the gene expression differences driving the emergence of each cell cluster, researchers can gain insights into key regulatory genes and pathways involved in cell fate decisions, development, and functional specialization. Based on the tree-structured diagram, CellScope introduces a novel multilevel gene identity system, referred to as dynamic ‘molecular identity.’ By analyzing the expression differences of genes among different cell clusters within the hierarchical levels of clustering, CellScope classifies genes into distinct identities, including housekeeping genes, moderately cell-type-related genes, and strongly cell-type-related genes **(Figure 1d** and **Methods E)**. By evaluating changes in gene identities across these clustering hierarchies, CellScope transcends the traditional binary classification of genes as either marker or non-marker genes.

### CellScope demonstrates superior performance in cell clustering and gene selection

We evaluated CellScope’s performance in cell clustering using 36 distinct scRNA-seq datasets with known cell types (**Supplementary Table S1, S2**). These datasets cover various human and mouse tissues—including brain, pancreas, embryos, and immune cells—and range widely in size (90 to 265,767 cells) and complexity (3 to 20 cell classes). Each dataset includes gold-standard cell type labels determined through methods like cell morphology and marker gene expression. We compared CellScope to two widely used single-cell analysis methods: Seurat [31] and Scanpy [12]. True cell-type labels was used only for post-hoc evaluation.

CellScope achieves the best cell clustering performance across all data sets regarding accuracy, robustness, and computational efficiency. To quantify clustering performance, we used multiple clustering evaluation metrics: adjusted rand index (ARI) [32], the clustering accuracy (ACC) [33], the normalized mutual information (NMI) [34], and jaccard index (JI) [35], where higher values indicate better clustering. **Figure 2a** shows ARI values for each dataset, along with mean and standard deviation. CellScope significantly outperformed the other methods, achieving the highest average ARI of 0.88 with the lowest standard deviation of 0.01. In comparison, Seurat had a mean ARI of 0.66 (standard deviation 0.03), and Scanpy had a mean ARI of 0.68 (standard deviation 0.04). CellScope ranked first in 32 out of 36 datasets and second in 3 datasets. The Wilcoxon signed-rank test confirmed CellScope’s superiority over Seurat (*p* = 2.91 × 10^−10^) and Scanpy (*p* = 2.56 × 10^−9^). Similar results across other metrics demonstrate CellScope’s advantages (**Supplementary Figure S1 and Table S3**). Beyond its excellent performance, CellScope showed superior computational efficiency (**Figure 2b and Supplementary Table S4**), with an average runtime less than half of Scanpy’s and less than a quarter of Seurat’s.

**Figure 2.**
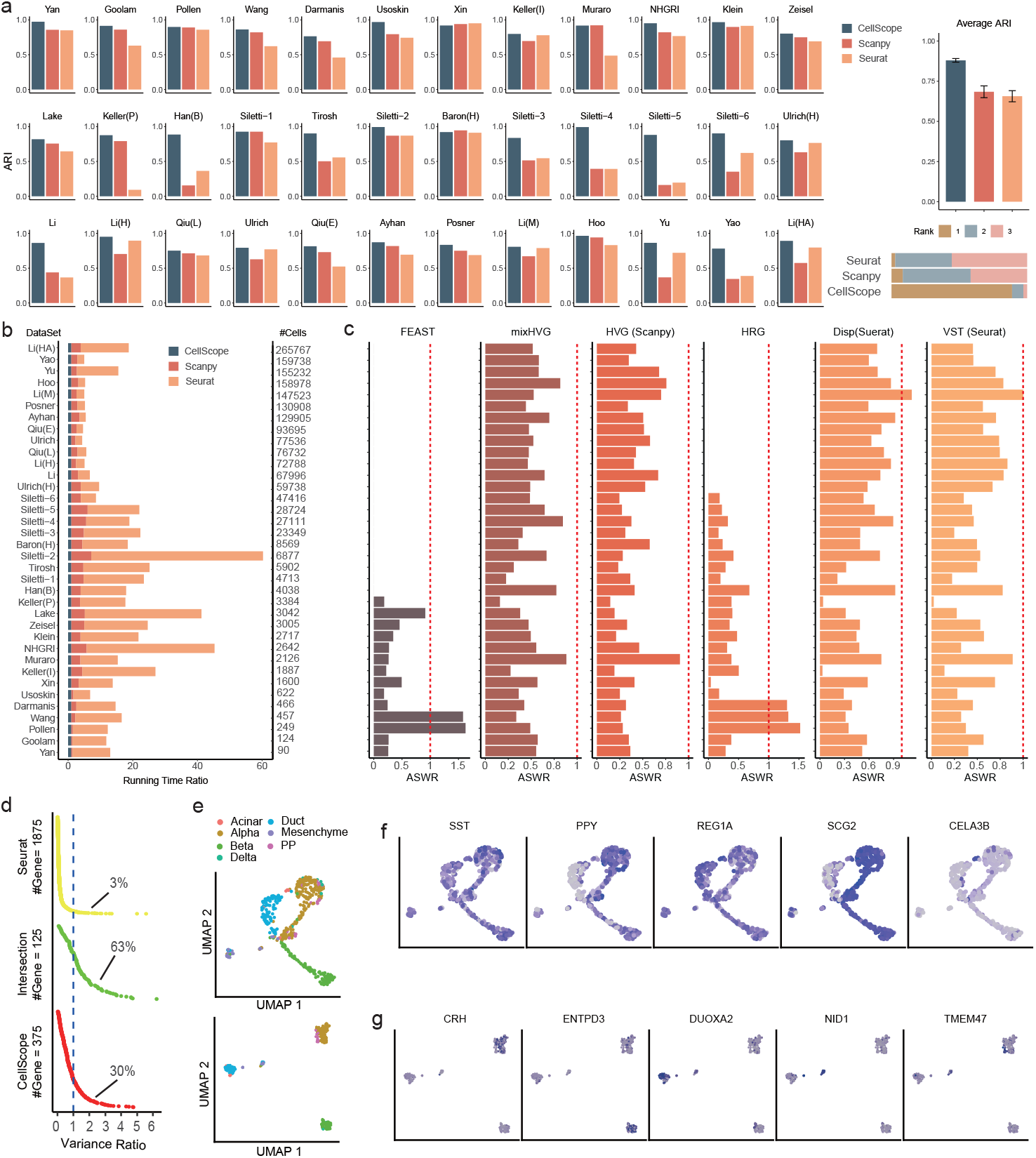
Performance of CellScope on 36 benchmark datasets. **a**. Clustering performance evaluation of CellScope, Seurat, and Scanpy using the Adjusted Rand Index (ARI). The first 36 panels display ARI values for individual datasets, while the final panel shows the mean ARI with standard deviation indicated by vertical bars. Bottom right shows the Rank distribution based on clustering performance. **b**. Execution time comparison across benchmark datasets. The x-axis depicts relative time ratios, with CellScope’s execution time set as the baseline of 1. Other methods’ execution times are expressed as multiples of CellScope’s time. **c**. Gene selection performance of CellScope compared to six gene selection methods across benchmark datasets. The x-axis depicts relative Average Silhouette Width Ratio(ASWR), with CellScope’s ASW set as the baseline of 1 (red dashed line). Other methods’ ASW are expressed as multiples of CellScope’s. Higher ASW indicates better performance. Absent bars indicate method failure or timeout. **d**. Gene Selection performance Comparison of CellScope and Seurat valued by variance ratio of Wang Dataset. CellScope can not only captures the high-quality marker genes identified by Seurat but also uncovers meaningful highly variable genes that Seurat overlooks. **e**. Visualization of CellScope (Bottom) and Seurat (Top) of human pancreatic cells from Wang et al. [40], colored by the true cell types. **f**. Visualization of the distribution of Marker genes selected by Seurat. Darker colors represent higher gene expression. It shows that many marker genes tend to be expressed across most cell types. **g**. Visualization of the distribution of Marker genes selected by CellScope.

Gene selection is crucial for clustering results. CellScope exhibits superior performance in this area compared to other methods, including Disp (Seurat) [36], VST (Seurat) [31], HVG (Scanpy) [12], mixHVG [37], FEAST [38], and HRG [39]. We use multiple evaluation metrics to assess the effectiveness of gene selection methods, including average silhouette width (ASW), variance ratio, cell-type local inverse simpson index (LISI), and KNN classification accuracy. These metrics collectively evaluate different aspects of gene selection effectiveness, including preservation of local structure preservation, cell type separation, cluster quality, and predictive power of the selected genes. Among these, ASW is highlighted in the main text due to its ability to provide a global view of clustering quality by simultaneously evaluating intra-cluster cohesion and inter-cluster separation. CellScope consistently achieves higher ASW values on the majority of datasets (**Figure 2c**), while other metrics, such as k-neighborhood purity and variance ratio, further confirm its superiority in gene selection effectiveness (**Supplementary Figure S4 and Tables S5-S9**).

**Figure 2e** compares the visualization of CellScope and Seurat using the human pancreatic cells from Wang et al. [40]. Seurat’s results incorrectly suggest that the three cell types—Alpha, Duct, and Beta—are interconnected and indistinguishable, whereas CellScope accurately achieves clear separation among the cell types. This difference is largely attributed to CellScope’s manifold fitting-based gene selection strategy. As shown in **Figure 2d**, only 125 genes overlap between Seurat’s 2000 and CellScope’s 500 selected genes, with 63% of these shared genes exhibiting significant variation (variance ratio *>* 1, defined as the ratio between a gene’s inter-cell-type and intra-cell-type expression variance, indicating its power in distinguishing cell types). More notably, 30% of CellScope-specific genes showed strong differential expression (variance ratio *>* 1) between cell classes, whereas only 3% of Seurat-specific genes exhibited such pronounced differences. This indicates that CellScope captures high-quality marker genes and discovers meaningful high-variance genes that Seurat overlooked. For instance, the expression patterns of Seurat-selected genes (e.g., *SST, REG1A*) show minimal cell-type specificity, resulting in poor cluster separation. In contrast, CellScope uniquely identifies genes (e.g., *CRH*) that are not selected by Seurat and exhibit highly specific expression patterns, facilitating more accurate cell type differentiation.(**Figure 2f-g**). A comparison between CellScope and Scanpy gene selection can be seen in **Supplementary Figure S2**, and results of other all benchmark datasets are available in **Supplementary Table S13-S15**.

### CellScope enhances the ability to distinguish similar cell types, detect rare types, and perform multi-level clustering

CellScope demonstrates its sensitivity to distinguish similar cell types and the the ability to detect rare cell populations. This was exemplified using the Human Brain cells dataset from NHGRI [41] and the Mouse Pancreas cells dataset from Keller(P) [42] (**Figure 3a,b**), we observed that CellScope’s visualization results showed clearer separation of cell types compared to popular methods like Scanpy and Seurat.

**Figure 3.**
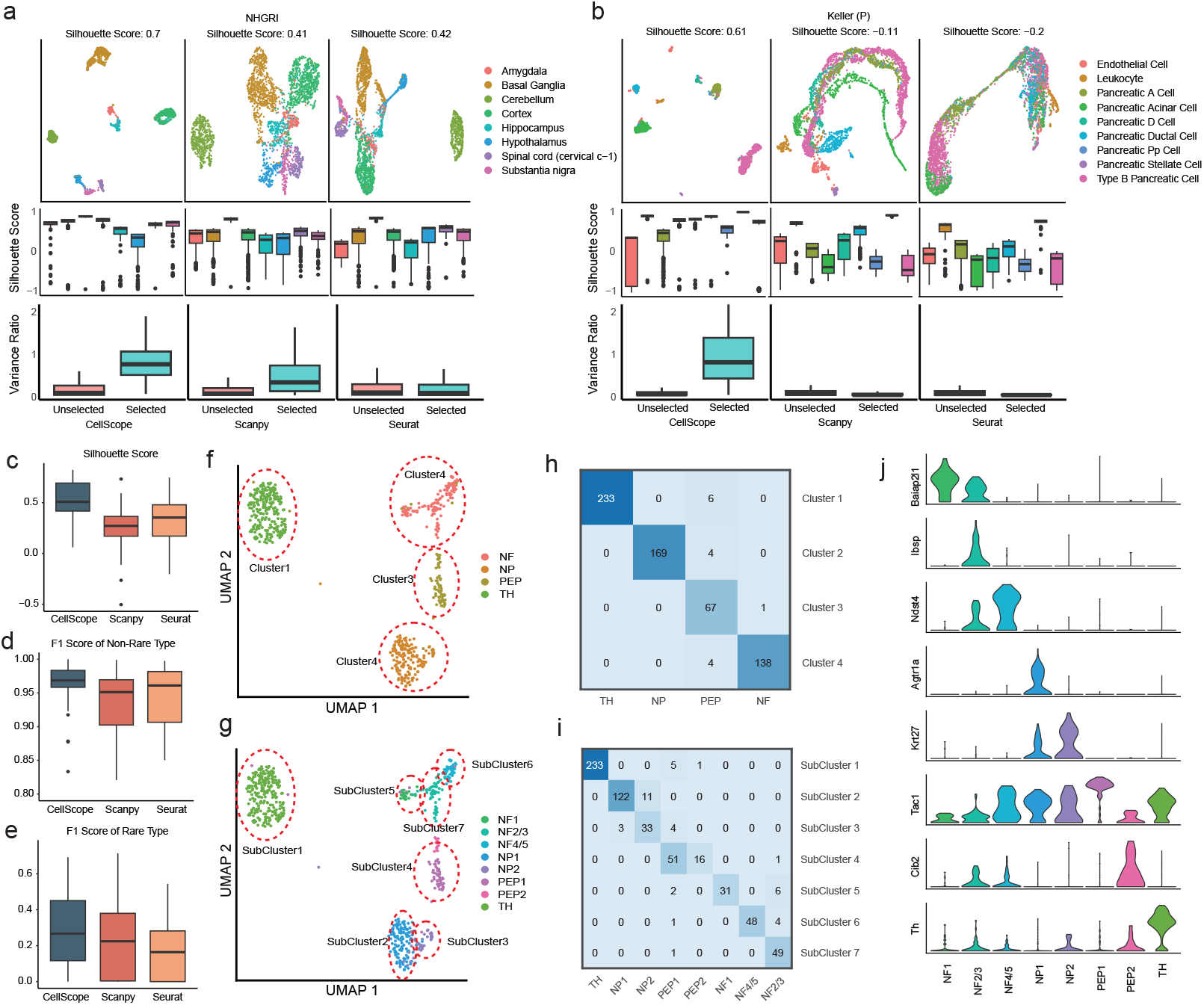
The performance of CellScope in distinguishing similar cell types, detecting rare types, and performing multi-level clustering: **a**. Upper panel: CellScope, Seurat, and Scanpy visualizations on the NHGRI dataset. Middle panel: Box plots of the Silhouette scores for each cell type. Lower panel: Box plots of the variance ratio for genes selected by the three methods compared to unselected genes. **b**. Upper panel: CellScope, Seurat, and Scanpy visualizations on the Keller (P) dataset. Middle panel: Box plots of the Silhouette scores for each cell type. Lower panel: Box plots of the variance ratio for genes selected by the three methods compared to unselected genes. **c**. Silhouette score-based visualization performance comparison across 36 datasets for CellScope, Seurat, and Scanpy. **d, e**. Cell-type recognition capability comparison. F1 scores across 36 datasets for non-rare (d) and rare (e) cell types using CellScope, Seurat, and Scanpy. **f**. CellScope visualization of the Usoskin dataset at cell type level. **g**. CellScope visualization of the Usoskin dataset at sub-cell type level. **h**. Confusion matrix between second-level clustering results and true cell types on the Usoskin dataset. **i**. Confusion matrix between fifth-level clustering results and true sub-cell types on the Usoskin dataset. **j**. Key marker genes identified by CellScope for the Usoskin dataset analysis.

Quantitative analysis revealed that for nearly all cell types in the NHGRI dataset, the Silhouette Score was significantly higher with CellScope compared to Scanpy or Seurat. Specifically, Scanpy and Seurat could only distinguish the Cerebellum from other categories, while the remaining cell types were mixed together without clear boundaries (Middle of **Figure 3a**). In contrast, CellScope’s results successfully separated Basal Ganglia, Cerebellum, and Cortex from other cell types. Moreover, Amygdala and Hippocampus clustered together as one group, and Hypothalamus, Spinal cord, and Substantia nigra formed another, with clear boundaries between each cell type within these groups (Upper of **Figure 3a**). This highlights the superiority of CellScope in distinguishing cell types.

The analysis of the Keller (P) dataset further demonstrated CellScope’s robust ability to recognize both common and rare cell types. For instance, common cell types like type B pancreatic cells, comprising approximately 40% of the total population, formed distinct clusters, while rare cell types, including leukocytes, pancreatic PP cells, and pancreatic stellate cells, representing 3.6%, 2.1%, and 1.4%, respectively, were distinctly separated. Compared to Scanpy and Seurat, CellScope’s advantage lies in its superior ability to identify cells across multiple scales. This capability can be attributed to our gene selection algorithm, which favors genes with higher inter- and intra-class variance ratios. This approach enhances the separation between classes and ensures that nearly every class’s markers are represented, even with only 500 selected genes (Lower of Figure 3a,b and Supplementary Figure S3).

Expanding this analysis to a comprehensive set of 36 datasets (**Supplementary Table S1**), CellScope consistently outperformed Scanpy and Seurat, with higher overall Silhouette Scores (**Figure 3c**). To further quantify its effectiveness in identifying both common and rare cell types, we defined non-rare cell types as those comprising more than 20% of the population, and rare types as those comprising less than 5%. For non-rare cell types, CellScope achieved high and stable F1 scores (**Figure 3d**), comparable to those of existing methods. However, for rare cell types, CellScope’s F1 scores were significantly higher (**Figure 3d**), underscoring its unique advantage in identifying low-abundance cell populations.

Additionally, CellScope demonstrated advanced multi-level clustering capabilities by analyzing the mouse lumbar cells from Usoskin dataset [43], which contains four major cell types and their eight subtypes. Using our hierarchical clustering, we first mapped major cell types at the second clustering level (**Figure 3f, h**), successfully separating tyrosine hydroxylase containing (TH), neurofilament containing (NF), peptidergic nociceptors (PEP), and non-peptidergic nociceptors (NP) populations. For subtypes without clear boundaries, we extended the clustering process (**Figure 3g, i**) to accurately identify NP and NF subtypes. This superior performance can be attributed to CellScope’s gene selection strategy, which identifies genes uniquely expressed in specific subtypes (**Figure 3j**). For instance, CellScope selects *Tac1* and *Th* as distinctive markers based on their specific expression in PEP1 and TH cells, respectively. Additionally, CellScope identified *Agtr1a* based on its unique expression in NP1 neurons. This gene selection approach, combined with multi-level clustering, enables accurate identification of cell types and subtypes while preserving the biological relationships between cell populations - a precision that Seurat and Scanpy could not achieve (**Supplementary Figure S4**).

### CellScope’s tree-structured visualizations refine characterization of brain cell atlases

CellScope’s tree-structured visualization effectively displays the hierarchical relationships between cell types and their subtypes. To demonstrate, we implemented CellScope with a dataset from the Human Brain Cell Atlas [44] named Siletti-1. First, CellScope’s tree-structured visualization helps capture all cell types of the Siletti-1 dataset. Specifically, CellScope categorized the red nucleus within the midbrain into nine distinct classes and identified nearly all superclusters previously reported in [44]. Notably, CellScope is able to identify the Fibroblast sub-type comprising mere 26 cells (**Figure 4a**).

**Figure 4.**
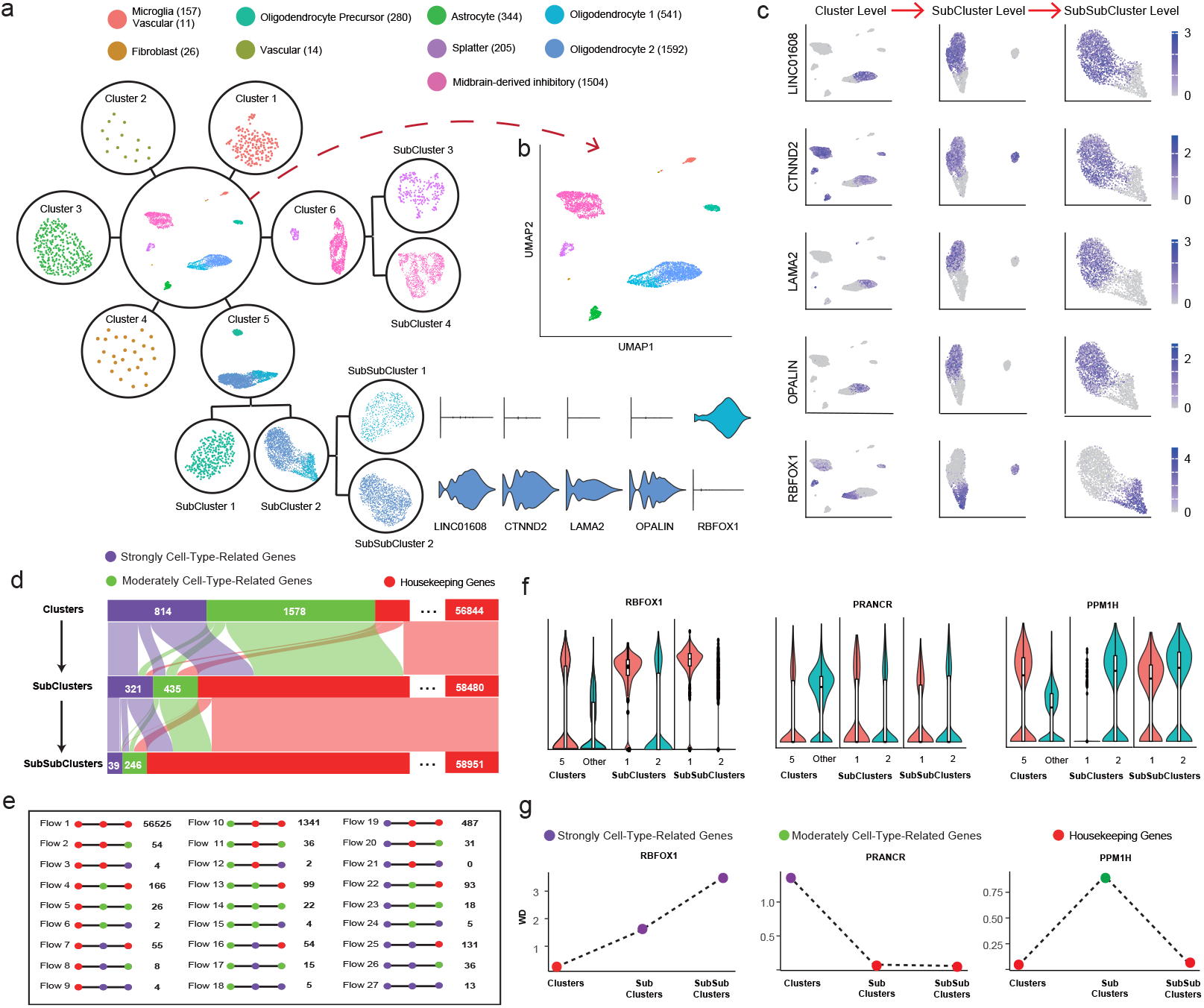
CellScope’s tree-structured visualizations enhance Human Brain Cell Atlas characterization. **a** CellScope tree-structured visualization displays novel cell types. **b** UMAP visualization of the whole dataset labeled by clusters identified by CellScope in Panel a. **c** Expression of selected marker genes (*LINC01608, CTNND2, LAMA2, OPALIN, RBFOX1*) shown at cluster, subcluster, and subsubcluster levels. **d** Sankey diagram of the proportions of three gene types (Strongly Cell-Type-Related Genes, Moderately Cell-Type-Related Genes, and Housekeeping Genes) in different levels of cell clusters. **e** Number of genes preserved in 27 gene type conversion flows among the three-level hierarchical cell clustering. **f** Violin plots of the expression of genes in three representative gene type conversion flows at different clustering levels. (From left to right: *RBFOX1* in Flow 6, *PRANCR* in Flow 19 and *PPM1H* in Flow 4). **g** Wasserstein distance (WD) of gene expression across three representative gene type conversion flows at different clustering levels. The distance is calculated for gene expression between Cluster 5 and the rest, SubCluster 1 and SubCluster 2, as well as SubSubCluster 1 and SubSubCluster 2. Smaller WD values indicate a similar gene expression level between compared clusters. (From left to right: *RBFOX1* in Flow 6, *PRANCR* in Flow 19, and *PPM1H* in Flow 4).

Second, our analysis revealed that Oligodendrocytes (OLs) can be further differentiated into two subtypes, tentatively designated as Oligodendrocyte1 (OL1) and Oligodendrocyte2 (OL2), containing 679 and 1454 cells, respectively. To elucidate the spatial and hierarchical expression patterns of key marker genes distinguishing these two subtypes, **Figure 4b, c** highlights the expression distributions of five differentially expressed genes across three clustering levels: Cluster, SubCluster, and SubSubCluster. Specifically, the high expression of the OL1 marker gene *RBFOX1* indicates that these cells are in a mature terminal state [44]. In contrast, the high expression of the OL2 marker gene *OPALIN* reflects active myelination, suggesting that OL2 cells are still in the differentiation stage. Meanwhile, the low expression of *OPALIN* in OL1 further supports the notion that OL1 cells have reached full maturity [45]. Additionally, *CTNND2* [46] plays a critical role in cell adhesion and synapse formation, while Laminin-2, encoded by *LAMA2* [47], regulates the spreading of oligodendrocytes (OLs) and myelination in the central nervous system via the integrin signaling pathway. Therefore, the cell population with high expression of these genes (OL2) likely interacts more closely with axons, further indicating that myelin structures are forming [48].

Third, CellScope’s multi-layer cell clustering provides novel perspective to study the dynamic role of marker genes. In other words, CellScope can further assigns each gene a new dynamic ‘molecular identity’ of marker genes depending on whether they are solely unique to one layer of clustering or they keep playing a marker’s role in all layers of clustering. In Siletti-1, CellScope demonstrates a three-level hierarchical clustering system, referred to as clusters, subclusters, and sub-subclusters, which progressively identifies homogeneous cell groups with increasing resolution (**Figure 4a**). By categorizing genes into three dynamic identities—housekeeping genes (HG), moderately cell-type-related genes (MCTRG), and strongly cell-type-related genes (SCTRG)—based on their significance across clustering levels (**Methods E** for details), we visualized the relationships between these gene identities using a Sankey diagram (**Figure 4d**). Additionally, the overlaps and transitions of these gene identities across the three clustering levels were analyzed (**Figure 4e**). As the clustering resolution increases, the number of Housekeeper Genes gradually increases, while the number of SCTRGs and MCTRGs gradually decreases. This phenomenon reflects the changing roles of genes at different clustering levels. Specifically, as the resolution increases, many genes transition from SCTRGs or MCTRGs to Housekeeper Genes. During cell differentiation, SCTRGs and MCTRGs are typically involved in establishing cell-specific functions or characteristics, especially during the early and middle stages of development. As cells progress into maturity, more Housekeeper Genes are activated, indicating that the cells have entered a phase focused on maintaining stable functions, such as protein synthesis, metabolism, and cell proliferation, which are essential for basic cell maintenance and biological processes.

We used three types of flow as examples (Flow 6 HG-SCTRG-SCTRG, Flow 4 HG-MCTRG-HG and Flow 19 SCTRG-HG-HG) and identified genes *RBFOX1* in Flow 6, *PPM1H* in Flow 4, and *PRANCR* in Flow 19, which serve as marker genes exclusively at the SubSubCluster, SubCluster, and Cluster levels, respectively (see **Figure 4f, g**). Specifically, in Flow 6, *RBFOX1* is widely expressed in the nervous system and regulates various alternative splicing events related to neural development and maturation, including transcription factors, splicing factors, and synaptic proteins [49]. *RBFOX1* shows high expression levels in both oligodendrocytes (OLs) and Splatter-type cells, while the expression difference between oligodendrocyte precursor cells (OPCs) and oligodendrocytes is minimal. This is likely because OPCs are the direct precursor cells of OLs, during which *RBFOX1* primarily supports basic cell development, differentiation, and gene regulation [50], maintaining developmental continuity. And as mentioned earlier, there is a significant expression difference between the two OL subtypes, OL1 and OL2, due to their different maturity levels.

In Flow 4, gene enrichment analysis using ClusterProfiler [51] reveals that *PPM1H* is enriched in functions related to dephosphorylation and protein regulation. During OL differentiation, *PPM1H* regulates specific protein dephosphorylation events, contributing to the synthesis and formation of myelin proteins. Interestingly, the expression differences of *PPM1H* between different OL subtypes are minimal, likely because mature OLs need to maintain a stable signaling environment for myelin formation, which results in consistent expression of *PPM1H* across different OL subtypes. In Flow 19, *PRANCR*, a long noncoding RNA known to regulate keratinocyte proliferation and cell cycle progression [52], shows low expression in Cluster 5, particularly in OPCs and OLs. This is consistent with its established role in cell proliferation rather than differentiation. Since myelinating cells are specialized and no longer proliferating, the low expression of *PRANCR* in these cells aligns with their specialized function in myelination [53].

### CellScope improves analysis of disease-control cell atlases

Disease-control cell atlases are invaluable in modern medical research, offering critical insights into disease mechanisms, potential therapeutic targets, and novel diagnostic approaches by comparing the cellular composition and functional states of healthy and diseased individuals. In complex diseases like COVID-19, these atlases provide the potential opportunities to help uncover cellular changes during disease progression, track immune responses, and identify cell subtypes associated with disease severity. While existing analysis pipelines [54] have made significant contributions, there remains room for improvement in detecting such signals, particularly in distinguishing cell populations associated with disease states.

To better analyze the disease-control atlas, we developed a CellScope-based analytical pipeline (**Supplementary Figure S5**) and applied it to peripheral blood mononuclear cell(PBMC) data from healthy individuals and COVID-19 patients with varying disease severities [54]. Focusing on the monocyte-dendritic cell system, we successfully identified and isolated this system in step 10 of the CellScope analysis (**Figure 5a**). CellScope clearly distinguished classical monocytes, non-classical monocytes, and conventional dendritic cells (middle, left, and right clusters), outperforming traditional UMAP (**Figure 5b**). It also revealed a continuous differentiation trajectory from classical monocytes to conventional dendritic cells and non-classical monocytes.

**Figure 5.**
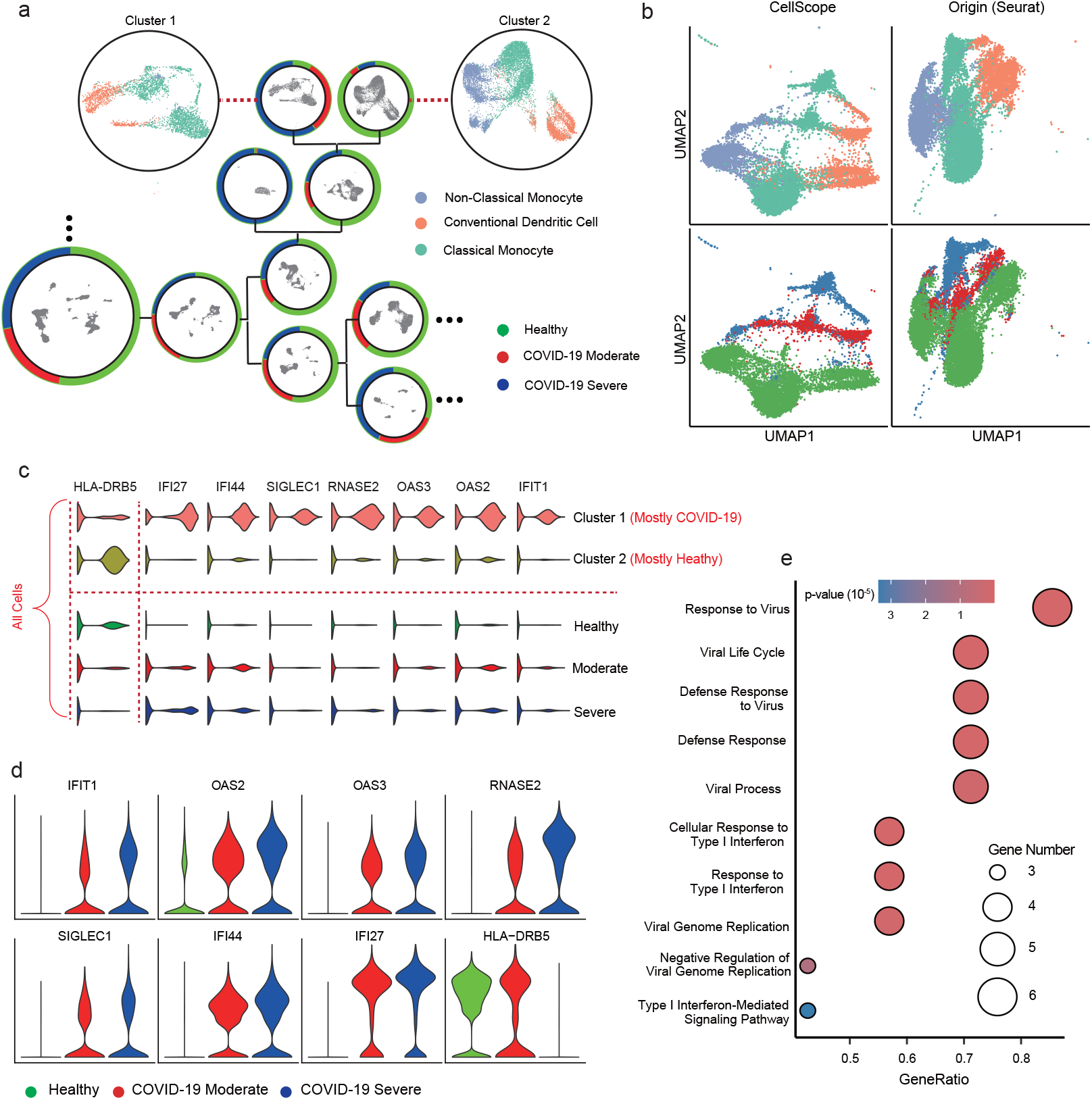
Tree-structured visualization, differential gene expression analysis, and GO enrichment for the COVID-19 PBMC dataset [54]. **a** Tree-structured visualization generated by CellScope. The colors inside the circle represent different cell types, while the outer ring colors indicate disease status. **b** Visualization comparison of the monocyte-dendritic cell system provided by CellScope and Origin (Seurat), with colors representing cell type and disease severity. **c** Violin plots illustrate the expression levels of 8 marker genes, arranged from top to bottom as Cluster 1, Cluster 2, and cells outside Cluster 1 and Cluster 2, categorized by healthy state, COVID-19 moderate, and COVID-19 severe. **d** Violin plots showing the expression levels of 8 marker genes in the monocyte-dendritic cell system across three disease states. **e** GO enrichment comparison of gene clusters uniquely upregulated in Cluster 1.

Moreover, CellScope, compared to Seurat, provides a clearer distinction between the three disease states—severe, moderate, and healthy. Specifically, in step 11, CellScope further refined the separation of COVID-19-associated cells, demonstrating its exceptional capability in identifying disease states. **Figure 5c** illustrates the expression of eight marker genes in Cluster 1 (mostly COVID-19) and Cluster 2 (mostly healthy). Among them, seven marker genes—*IFIT1, OAS2, OAS3, RNASE2, SIGLEC1, IFI44*, and *IFI27*—exhibit significantly higher expression in Cluster 1, while *HLA-DRB5* shows elevated expression in Cluster 2, providing key molecular markers for distinguishing disease states. The expression levels of these genes in the monocyte-dendritic cell system increase progressively with disease severity (**Figure 5d**), highlighting the marked differential expression between COVID-19 and healthy states. Notably, such differences were not observed in other cell types (**Figure 5c**). This likely underscores the critical role of the monocyte-dendritic cell system in viral recognition, antiviral responses, and immune regulation. The upregulation of Cluster 1 genes highlights a robust immune defense against SARS-CoV-2, particularly interferon-mediated antiviral responses, which are crucial components of the innate immune system’s defense against viral pathogens [55]. This gene expression pattern not only indicates active viral infection but also reveals the complex interactions between the virus and the host immune system.

SARS-CoV-2 appears to impair dendritic cell function by downregulating *HLA-DRB5* expression, leading to a loss of antigen-presentation capacity in infected monocytes and dendritic cells, facilitating viral evasion of T cell-mediated immune responses [56]. Additionally, clusterProfiler [51] enrichment analysis of the seven genes highly expressed in Cluster 1 (**Figure 5e**) revealed their key roles in antiviral immune responses. *IFIT1* and *IFI27* are strongly linked to interferon responses, while *OAS2, IFIT1, and SIGLEC1* activate interferon signaling, triggering antiviral defenses and inhibiting viral replication. These genes also regulate viral RNA replication, halting virus proliferation. The enrichment analysis further uncovered their roles in multiple stages of the viral lifecycle, from recognition to response and clearance, highlighting their importance in antiviral immunity.

### CellScope demonstrates interpretability, robustness, and user-friendliness

CellScope demonstrates significant advantages in algorithm interpretability, robustness, and user-friendliness. It efficiently identifies biologically meaningful key genes, adapts seamlessly to diverse datasets, and offers intuitive and informative visualization. In contrast, existing tools like Seurat and Scanpy primarily focus on highly variable genes for gene selection, which may face certain limitations in distinguishing between housekeeping genes and key genes. These tools also often require careful parameter adjustment to achieve optimal results.

First, CellScope enhances algorithm interpretability through its innovative gene selection approach. The method selects samples with high density and large distances from each other as “manifold seeds” to effectively distinguish between noise and meaningful signal space. This approach is grounded in a fundamental principle of unsupervised learning and cluster analysis—the relationship between local density and distance on manifolds [57]. The intuition behind this strategy is that high-density points have a greater probability of residing at the centers of manifold clusters, where their local neighborhoods typically exhibit higher purity in terms of class composition. Analysis of the human brain cells from Darmanis dataset [58] revealed a strong negative correlation between local density and distance from true cluster centers (**Figure 6a**), where high-density points consistently exhibited improved neighborhood purity (**Figure 6b**), supporting our density-distance based manifold seeds identification method. We also compared the differences between selected and unselected genes within and between clusters (**Figures 6c, d**). Compared to the unselected genes, the genes selected by CellScope exhibit significantly larger inter-class differences and smaller intraclass differences (*p* = 3.8 × 10^−38^). In contrast, the unselected genes show that intra-class variance is significantly greater than inter-class variance. This enhanced distinction facilitates better cluster separation, demonstrating that our gene selection process effectively captures the most informative genes for distinguishing different cell types.

**Figure 6.**
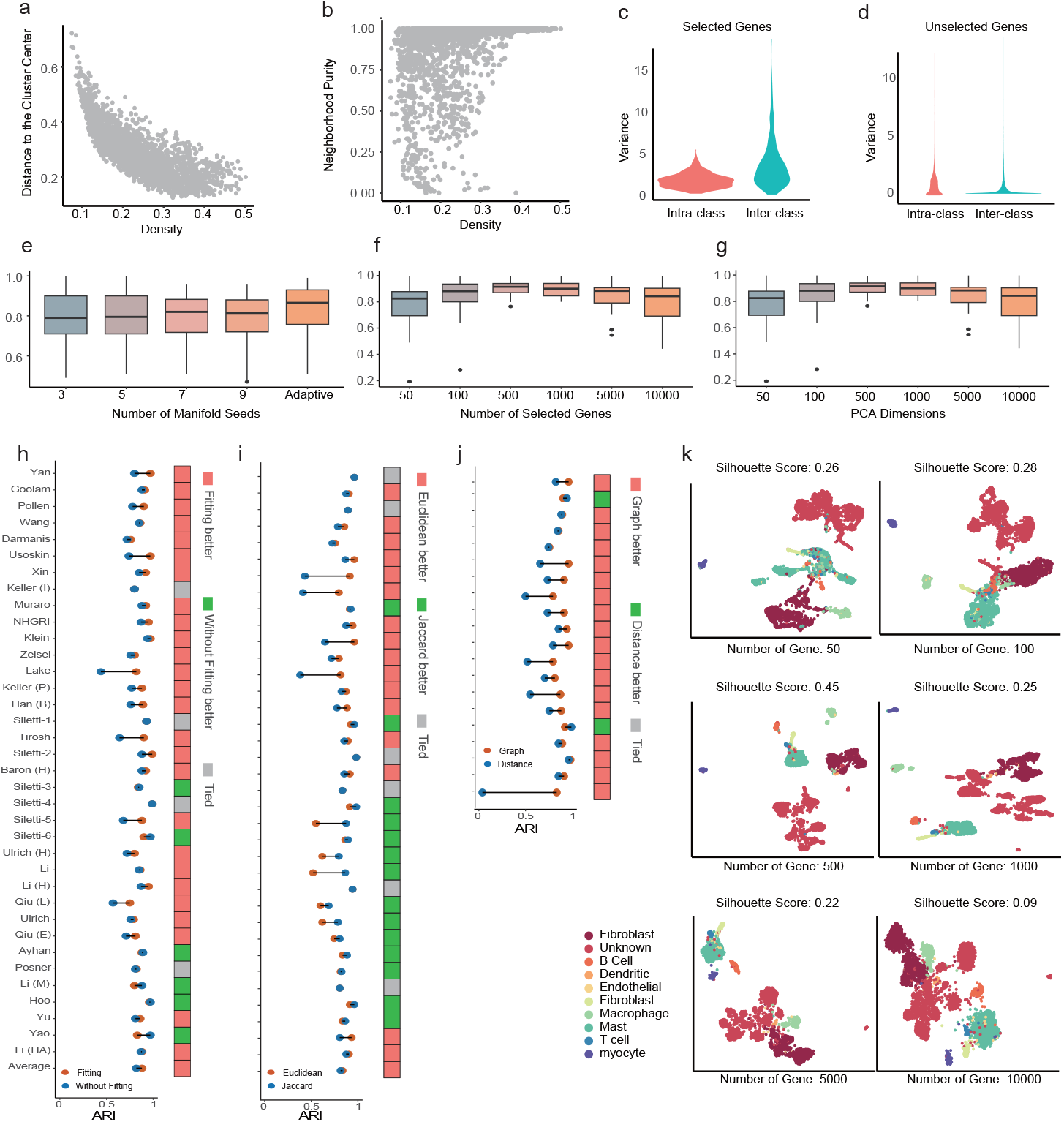
CellScope demonstrates interim results optimal parameter choices with a strong theoretical foundation. **a** Scatter plot of Density vs Distance for Darmanis dataset. The X-axis shows the local density of every cell. The Y-axis shows the distance from the cell to its nearest true class center. **b** Scatter plot of Purity vs Distance for Darmanis dataset. The X-axis shows the 100-nearest neighbors of every cell. The Y-axis shows the distance from the cell to its nearest true class center. **c** Violin plot comparing intra-class and inter-class gene variance for selected genes using the CellScope method in the NHGRI dataset. **d** Violin plot comparing intra-class and inter-class gene variance for unselected genes using the CellScope method in the NHGRI dataset. **e** Box plot of ARI values with different quantities of manifold seeds in the gene selection part of CellScope across small to medium-scale datasets with less than 50,000 cells. **f** Box plot of ARI values with different numbers of selected genes in the gene selection part of CellScope across small to medium-scale datasets with less than 50,000 cells. **g** Box plot of ARI values with different dimensions of PCA in the gene selection part of CellScope across small to medium-scale datasets with less than 50,000 cells. **h** Visualization of ARI values comparison with methods on whether the manifold is fitted. The Orange above shows that the Manifold Fitting method has better performance for almost all datasets. **i** Visualization of ARI values comparison with different clustering metrics. The Orange above shows that the Euclidean method has better performance when dataset sizes are small. The green squares above indicate that Jaccard-based method performs better when dataset sizes are large. **j** Visualization of ARI values comparison with different hierarchies. The Orange above shows that the Graph-based method has better performance than distance-based methods across small to medium-scale datasets with less than 25,000 cells. **k** Cluster result visualization with Silhouette Scores of Tirosh dataset using different number of genes selected in Gene Selection part of CellScope.(from left to right: 50, 100, 500, 1000, 5000, 10000).

Second, CellScope demonstrates exceptional robustness and adaptability by minimizing manual parameter tuning and maintaining consistent performance across various datasets. Unlike existing algorithms that often require meticulous manual parameter tuning, CellScope offers robust performance and ease of use across various settings.We performed comprehensive robustness analyses to evaluate key parameters in our gene selection process, including the dimensions of PCA, the quantity of selected manifold seeds, and the number of selected genes. Firstly, we assessed how varying the dimensions of PCA affects clustering performance. The ARI remained stable across different dimensions of PCA (**Figure 6g**), with optimal performance observed at around 100 dimensions. This stability indicates that CellScope is not overly sensitive to the choice of dimensions of PCA. Secondly, we compared our adaptive method for manifold seeds selection with fixed-number approaches. Our adaptive method consistently outperformed fixed-number methods across different dataset sizes (**Figure 6e**). This adaptability allows CellScope to automatically adjust to the characteristics of each dataset without manual intervention. Thirdly, we investigated the impact of the number of genes selected during gene selection. The clustering performance remained consistent over a wide range of gene counts (**Figure 6f**), with optimal results achieved around 500 genes. However, selecting too few genes (e.g., 50 genes) led to insufficient clustering information due to the omission of key genes determining cell identity, resulting in category confusion. Conversely, selecting too many genes (e.g., 10,000 genes) introduced redundant information, which masked cell type-specific signals and hindered effective separation of different cell types. This phenomenon was evident in the analysis of the human oral cavity cells of Tirosh dataset [59] (**Figure 6k**). These results emphasize the importance of judicious gene selection in single-cell analysis and highlight CellScope’s robustness and adaptability in handling this critical parameter.

Finally, CellScope prioritizes user-friendliness through several key design choices that optimize clustering analysis and enhance adaptability. To evaluate the importance and necessity of these key components in CellScope, we conducted a comprehensive evaluation of its clustering approach. We first investigated the impact of the cell projection component in the manifold fitting process on CellScope’s performance. **Figure 6h** illustrates that, for the majority of datasets, the application of manifold fitting yielded superior results compared to its absence. We then compared the performance of the graph-based hierarchical clustering method adopted in CellScope with traditional distance matrix-based hierarchical clustering. The ARI values showed that our graph-based clustering consistently outperformed distance-based clustering across almost all datasets examined (**Figure 6j**). This superior performance can be attributed to the ability of graph-based algorithms to more effectively capture local structures and nonlinear relationships in the data by constructing nodes and edges. This is particularly beneficial in high-dimensional datasets and clusters with complex morphologies, where traditional distance-based algorithms may struggle due to their inability to capture intricate structural properties. To further optimize CellScope, we implemented an adaptive distance metric based on dataset size: Euclidean distance for smaller datasets and Jaccard distance for larger datasets. The results (**Figure 6i**) confirm the reliability of this adaptive strategy. The effectiveness of this approach stems from the fact that Euclidean distance can introduce dimension-re/.lated artifacts in large datasets, a phenomenon known as the “curse of dimensionality” [60]. In contrast, the Jaccard distance metric effectively alleviates this issue by focusing on the presence or absence of features rather than their magnitude, making it more suitable for high-dimensional data.

## Discussion

In order to address fundamental challenges in single-cell RNA sequencing analysis, including biased gene selection, oversimplified cellular visualization, and limited capability in discovering and interpreting complex biological phenomena, we present CellScope, a comprehensive computational framework built upon manifold learning principles. CellScope introduces three key innovations: a rapid and accurate gene selection method that minimizes bias while maintaining biological relevance, a tree-structured visualization framework that comprehensively represents cellular hierarchies, and a multi-level characterization system that provides dynamic gene classifications across different resolutions. Based on these innovations, CellScope demonstrates comprehensive superiority: not only achieving exceptional accuracy, computational efficiency, and interpretability across diverse datasets, but also exhibiting powerful capabilities in discovering cell subpopulations and identifying disease-specific clusters.

A particularly powerful aspect of CellScope is its gene selection methodology, which demonstrates marked improvements over existing approaches. Current single-cell gene selection frameworks exist in two extreme states: overly simplistic or excessively complex. Simplistic algorithms like HVG, implemented in Seurat or Scanpy, directly select highly variable genes without distinguishing whether the variations arise from biological signals or noise. In contrast, complex algorithms typically employ pre-clustering or consensus approaches which, although performing better than HVG, are computationally intensive and can introduce biases from poor separation in the pre-clustering step. CellScope bridges this gap by introducing a balanced and efficient approach. By constructing reference clusters using only a small subset of trustworthy cells from the centers of high density regions, CellScope is able to identify biologically informative genes with higher reliability and efficiency than other established methods.

Popular visualization techniques like UMAP and t-SNE prioritize global structure preservation at the expense of local relationships, leading to a loss of fine-grained information about cell states and developmental trajectories. CellScope addresses this limitation through its innovative tree-structured visualization framework, which provides a hierarchical representation of cellular relationships across multiple resolutions. Unlike traditional dimensionality reduction methods that compress all information into a single view, our tree structure preserves both broad cellular categories and subtle cell states, enabling researchers to explore cellular hierarchies at different levels of granularity. This multi-resolution visualization approach is particularly powerful when analyzing complex tissues or disease progressions, where cellular states exist along continuous spectrums rather than discrete categories.

An important innovation of CellScope is its introduction of a multi-level identity system for genes, extending beyond traditional binary classifications of marker versus non-marker genes. By characterizing genes through their roles across multiple clustering layers—as housekeeping genes (HG), Moderately Cell-Type-Related Genes (MCTRG), or Strongly Cell-Type-Related Genes (SCTRG)—we establish a dynamic “molecular identity” for each gene. This hierarchical gene identity system reveals how genes can play different roles at different levels of cellular organization, providing crucial insights into the context-dependent nature of gene function. This understanding is particularly valuable for disease studies, where genes may acquire new functions in pathological states, and for developmental biology, where genes often switch roles during different stages of cellular differentiation.

Furthermore, CellScope demonstrates several compelling advantages in the analysis of single-cell data through its robust, user-friendly, and interpretable framework. The tool’s interpretability is grounded in its theoretically sound approach to manifold fitting [61], allowing it to effectively distinguish signal from noise. In addition, unlike other methods, CellScope does not require extensive parameter tuning, as evidenced by its stability across various parameter settings, including PCA dimensions, gene selection counts, and manifold seed selection methods. In conjunction, CellScope’s analytical rigor and practical usability position it as a reliable and accessible tool for single-cell analysis.

Benefiting from these technical advances, CellScope enables significant biological discoveries across diverse applications. For instance, our analysis of the Human Brain Cell Atlas uncovered two previously unrecognized oligodendrocyte subtypes which exhibit distinct molecular signatures characterized by *RBFOX1* and *OPALIN* expression, respectively, revealing new insights into myelination processes. CellScope’s capabilities also extend to disease research, as demonstrated in our COVID-19 study. Through analysis of PBMCs from patients with varying disease severities, we successfully distinguished traditional immune cell types while simultaneously identifying disease-specific states. The method revealed eight marker genes showing progressive expression changes with disease severity, specifically within the monocyte-dendritic cell system, providing crucial insights into antiviral immune responses. These discoveries in both steady-state and disease contexts demonstrate CellScope’s unique power in revealing both subtle cellular states and disease-associated transitions that are often missed by conventional analysis methods. Overall, CellScope provides a user-friendly, technically sound tool for advancing the field of single-cell omics in an era of increasing availability of large and complex single-cell datasets.

## Methods

CellScope aims to analyze single-cell data through manifold fitting, enabling precise identification of differences between cell types and subtypes. By identifying highly reliable cliques, the method effectively selects type-determined genes with class-specific differences while leveraging manifold fitting to mitigate the impact of technical noise. CellScope then constructs a cell-to-cell similarity graph and performs agglomerative clustering based on this graph to generate a hierarchical structure of cells. A tree-structured visualization intuitively represents the hierarchical relationships and reveals differentiation pathways among cells. Furthermore, CellScope analyzes gene expression changes along differentiation pathways and expression differences within the same hierarchy, providing new insights into gene functionality.

The CellScope workflow consists of five main steps, as explained in detail below: (A) Manifold fitting step 1, (B) Manifold fitting step 2, (C) Graph-based agglomerative clustering, (D) Tree-structured visualization, and (E) Characterization of genes from different categories.

We summarize the main notations used in the description of the method as follows: suppose we have a expression counts matrix 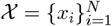, where each vector *x*_*i*_ corresponds to the expression values 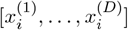 of the *i*-th cell across *D* genes. We also let 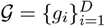, where *𝒢* represents the collection of genes related to *𝒳*. After manifold fitting step 1, we obtain 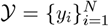, where *𝒴* represents the scRNA-seq data after manifold fitting step 1. Each vector *y*_*i*_ corresponds to the expression values 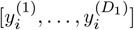 of the *i*-th cell across *D*_1_ selected genes, where *D*_1_ ≪ *D*. The set of selected genes is denoted as 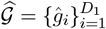, where 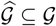. Subsequently, after manifold fitting step 2, we obtain 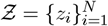, where *𝒵* represents the fitted scRNA-seq data for subsequent downstream analysis.

### A. Manifold fitting step 1

#### A1. Data preprocessing

We applied consistent preprocessing to all single-cell RNA sequencing datasets. First, log normalization (base 2) was applied to the raw data *𝒳*. Then, we normalized each cell, which is a standard procedure prior to downstream analyses such as principal component analysis (PCA). This step is essential for eliminating differences in total expression levels between cells, which may arise from technical or biological factors. It ensures that highly expressed genes in cells with elevated overall expression do not dominate the dimensionality reduction process, thereby preventing bias in subsequent analyses. Let 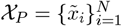 denote the preprocessed data, where each vector 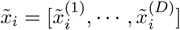 represents the expression values of the *i*-th cell after preprocessing. The set of genes after preprocessing is denoted as 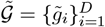.

### A2. Find highly reliable cliques

We begin this process with Principal Component Analysis (PCA), aiming to reduce noise and complexity in the data while retaining the primary sources of variation, thereby clarifying the manifold structure of the data and providing a solid foundation for subsequent manifold exploration. In PCA, we set the target dimensionality to *n*_1_ (defaulting to 100) and apply it to the preprocessed data *𝒳*_*P*_. This results in a collection of cells represented in a low-dimensional space, denoted as 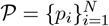, where 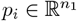.

We aim to identify highly reliable cliques associated with distinct submanifolds, beginning with the identification of the centers of these submanifolds, referred to as manifold seeds. Inspired by [57], local density reflects the compactness of a cell’s surrounding distribution, while relative distance measures the separation of a cell from other cells with higher density. Manifold seeds tend to exhibit significantly higher values in both metrics. Therefore, we evaluate the potential of each cell *p*_*i*_ to serve as a manifold seed by calculating its local density *ρ*(*p*_*i*_) and relative distance *δ*(*p*_*i*_), defined as:

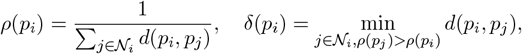

where *𝒩*_*i*_ is the set of *k* nearest neighbors of cell *p*_*i*_ (with *k* = 20 by default), and *d*(·, ·) denotes the distance between two cells, defaulting to Euclidean distance. Next, we compute the composite metric:

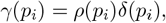

and select the cells with the highest *γ*(*p*_*i*_) values as manifold seeds. Specifically, we choose the top *m* cells with the largest *γ*(*p*_*i*_) values as manifold seeds, with *m* being a predefined hyperparameter which refers to the number of submanifolds hypothesized in advance.

However, since *m* is related to the number of cell types, an intelligent approach is required to determine the number of manifold seeds *m* when the number of cell types is unknown. We first address the issue of scale differences between *ρ* and *δ* in the calculation of *γ*. Both *ρ* and *δ* are normalized as

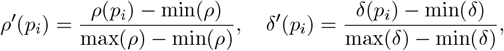

where 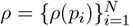 and 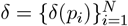. Next, we compute a scaleless index by multiplying the two normalized metrics:

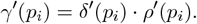

We then sort 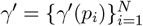 in descending order, with 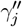 representing the *j*-th value in the sorted list. To further refine the selection of manifold seeds, we introduce the relative rate of change in *γ*^′′^:

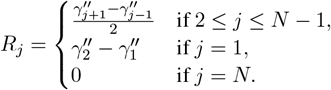

Finally, we select the manifold seeds that satisfy the following conditions:

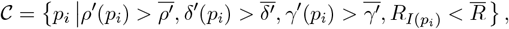

where 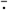 denotes the mean value of the respective set and 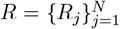. The index *I*(*p*_*i*_) refers to the position of *γ*^′^(*p*_*i*_) in *γ*^′′^ after sorting *γ*^′^. The combined metric-based strategy ensures that the selected manifold seeds possess both high local density and relative distance. The introduction of the relative rate of change further optimizes this selection process, intelligently selecting all high-confidence manifold seeds. Ultimately, we denote the set of selected seeds as *C* = {*c*_1_, · · ·, *c*_*m*_}, where *m* represents the number of seeds selected.

Given that the identified manifold seeds are highly likely to reside at the centers of their respective submanifolds, the cells in the immediate neighborhood of each seed are assumed to belong to the same class as the seed. Therefore, for each seed *c*_*i*_, we select its *k*_1_ nearest neighboring cells (with *k*_1_ = 5 by default), denoted as 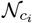. To ensure a high-confidence classification of the sets 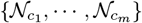, we combine the concept of connected components in graphs, define the following partition:

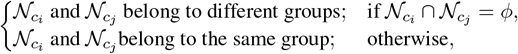

for 1 ≤ *i≠ j* ≤ *m*. The resulting partition of high reliable cliques 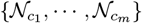 is recorded as *𝓁*_1_.

#### A3. Signal space identification

Due to the significant variance differences of genes in the signal space both within and between cell clusters, our goal was to identify genes that exhibit notable expression differences, particularly between different cell types, within highly reliable cliques. To achieve this, we leveraged the high-confidence labels *𝓁*_1_ obtained from these reliable cliques and performed a one-way analysis of variance(ANOVA [62]) on each preprocessed gene 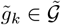. For each gene, the corresponding p-value was computed based on its expression across different cell clusters. We then selected the top *D*_1_ genes with the lowest p-values (default: *D*_1_ = 500), denoted as 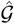, as the result of gene selection. Furthermore, we retained the genes from the signal space in *𝒳*_*P*_, denoted as *𝒴*.

These selected genes represent those with the most significant expression differences between submanifolds and are considered key to capturing the biological distinctions between different cell types. The set of genes in signal space,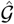, provides a group of genes that best distinguish the identified cell clusters, facilitating more efficient and biologically meaningful downstream analyses.

### B. Manifold fitting step 2

To further highlight the true biological signals and better reflect the underlying cellular heterogeneity, we project cells located between submanifolds or near manifold boundaries closer to the centers of their respective submanifolds. This process ensures that the boundaries between submanifolds become clearer after projection. We assume that the density of data points decreases as the distance from the manifold center increases. Therefore, our fitting process focuses on low-density points, as they are more likely to be influenced by noise.

First, we calculate the local density *ρ*(*y*_*i*_) for each cell *y*_*i*_ to assess its position within the manifold, using the following formula:

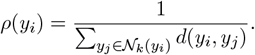

We then select the 5% of cells with the lowest densities to form the set of manifold outliers *𝒪*.

We assume that the closest high-density point to each outlier belongs to the same class and is closer to the center of its respective manifold. To achieve this, we adopt the projection estimation method from our previous work [25]. For each outlier *y*_*i*_ ∈ *𝒪*, the nearest high-density point *ŷ*_*i*_ is defined as:

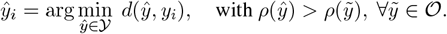

Subsequently, while preserving the relative position between the outlier *y*_*i*_ and *ŷ*_*i*_, we project *y*_*i*_ closer to the center of the manifold, as given by:

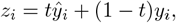

where *t* is 0.9 by default. For regular points not identified as outliers, we define their projection as *z*_*i*_ = *y*_*i*_, if *y*_*i*_ ∉ *𝒪*. The final dataset after the second step of manifold fitting is represented as 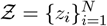.

### C. Graph-based agglomerative clustering

In the realm of unsupervised learning, graph-based clustering aggregation techniques are considered highly effective. A similarity graph can capture the local neighborhood relationships between points within a submanifold, reflecting the manifold’s local geometric properties. At the same time, through the construction of neighborhood relationships, the graph can represent the connectivity between different submanifolds. Therefore, after obtaining a clear manifold structure, we begin by using the Uniform Manifold Approximation and Projection (UMAP) algorithm [30] to construct the similarity matrix *S*. Specifically, for each cell *x*_*i*_, we determine its *k* = ⌈ log_2_(*N*) ⌉ nearest neighboring cells. We first set the similarity between the cell and cells outside its k-neighborhood to 0. Next, using a Gaussian kernel function, we compute the local similarity *s*_*ij*_ between two cells *z*_*i*_ and *z*_*j*_:

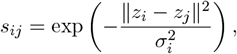

where *z*_*j*_ is in the *k*-neighborhood of *z*_*i*_ and ∥*z*_*i*_ − *z*_*j*_∥ denotes the Euclidean distance, and *σ*_*i*_ is a locally adaptive scale parameter satisfying:

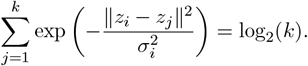

To construct a symmetric similarity graph, we define the edge similarity *w*_*ij*_ as:

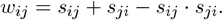

The corresponding symmetric similarity matrix is denoted as *W* = {*w*_*ij*_}. Finally, using the average linkage method based on pairwise similarities, we systematically merge data points starting with each cell as a separate cluster, and denote the clustering results after *K* steps as *T*_*K*_. *T*_*K*+1_ merges the two clusters with the highest average similarity, i.e., clusters *A*_*𝓁*_ and *B*_*𝓁*_ that maximize:

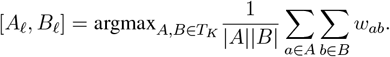

For exceptionally large datasets (over 100,000 data points), we reduce computational intensity by randomly selecting a subset of data (defaulting to 100,000 points), denoted as *S*_1_ and *S*_2_ for selected and unselected cells respectively. We recompute the adjacency matrix *W*_1_ for subset *S*_1_.

Using the same average linkage method, we obtain clustering results 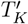 for the subsampled cell set *S*_1_. For cells in set *U*, we assign each cell *z* ∈ *U* to a cluster *C*_*𝓁*_ based on maximum average similarity to clusters in 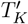:

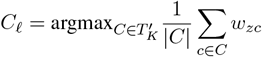

### D. Tree-structured visualization

To comprehensively and systematically explore cell types and their subtypes, we developed a tree-structured visualization method. First, we applied the UMAP algorithm [30] to visualize all cells and used the clustering results *T*_*N*_ − _1_ as the root node. Then, each cluster from *T*_*N*_ − _1_ was visualized individually to generate the first layer of child nodes. By comparing the clustering results of *T*_*N*_ − _2_ and *T*_*N*_ − _1_, we identified two new subtypes within the cells of the first-layer child nodes and visualized them as the second layer of child nodes. This iterative process was repeated in the same manner, advancing layer by layer.

To clearly illustrate the distribution of each pair of subtypes within their parent nodes, we applied a color-coding scheme to the tree-structured visualization. Specifically, each branch’s outermost nodes were colored first, and the colors of the child nodes were then propagated back to their respective parent nodes. This core visualization method effectively reveals the hierarchical relationships between cell types, illustrating how different cell populations emerge, differentiate, and specialize over time.

### E. Characterization of genes from different categories

After generating the Tree-structured visualization, we quantified the gene expression differences between sibling nodes (share the same direct parent node) using the Wasserstein distance. This metric measures the minimal transport cost required to transform one probability distribution into another, providing a robust comparison between gene expression profiles from sibling nodes.

For gene *g*_1_, assume its expression in two sibling nodes is represented by *P* = {*p*_1_, …, *p*_*n*_} and *Q* = {*q*_1_, …, *q*_*m*_}, respectively. First, both expression vectors are sorted in ascending order, yielding the ordered vectors 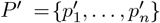 and 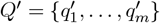. We then compute the cumulative distribution functions (CDFs) for the sorted vectors *P* ^′^ and *Q*^′^, as defined by the following equations

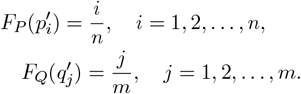

We then use linear interpolation to align the two CDFs *F*_*P*_ and *F*_*Q*_ onto the same probability space. For a given probability value *x* ∈ [0, 1], if 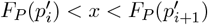, then 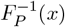) is calculated as

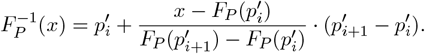

Similarly, 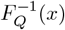 is computed in the same manner. Finally, the Wasserstein distance [63] is defined as the integral of the absolute difference between the two inverse CDFs across the probability space [0, 1]

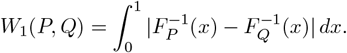

Finally, based on the Wasserstein distance calculated from the expression differences of genes between sibling nodes, we classified the genes into three categories

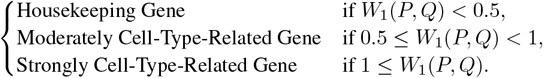

The threshold selection for Wasserstein distance is based on **Supplementary Figure S6**, using SubCluster1 and SubCluster2 in Siletti-1 **Figure 4** as an example to demonstrate its rationale. When the Wasserstein distance is less than 0.5, the gene expression distributions are similar. For distances between 0.5 and 1, there are notable differences in means, though some overlap remains, indicating moderate differences in gene expression. When the distance exceeds 1, the first quartile for the cluster with higher mean expression surpasses the third quartile of the other, indicating significant differences in gene expression.

### F. Benchmarks methods

#### F1. Compared pipelines

We compared CellScope with two existing popular baseline methods, Scanpy [12] and Seurat [31] in this study.Scanpy was implemented from its original source code repository (https://github.com/scverse/Scanpy). Highly variable genes were identified based on specified thresholds for mean expression and dispersion, and clustering was performed using the Leiden algorithm across a range of resolutions. The algorithm parameters were set according to the default parameter settings in the tutorial(https://Scanpy-tutorials.readthedocs.io/en/latest/pbmc3k.html).

Seurat was implemented from its source code (https://satijalab.org/seurat) with a scale factor of 10,000. We identified 2,000 variable features using the vst selection method as mentioned in Seurat tutorial(https://satijalab.org/seurat/articles/tutorial). Neighbors were identified using the first 10 principal components, and clustering by Louvain algorithm was performed across a range of resolutions.

In our experiments, we considered a range of clustering resolutions for both Scanpy and Seurat, specifically testing resolutions of 0.1, 0.15, 0.2, 0.3, 0.4, 0.5, 0.6, 0.7, 0.8, 0.9, 1.0, 1.1, 1.2, 1.3, 1.4, 1.5, 1.6, 1.7, 1.8, 1.9, and 2.0. For each method, we computed the Adjusted Rand Index (ARI) between the predicted clusters and the true cell type labels at each resolution. The final results for both methods were selected based on the highest ARI observed across all tested resolutions, ensuring that the best clustering performance was captured for each dataset.

#### F2. Gene selection methods

We compared several common gene filtering methods and the latest gene selection methods to identify the most effective techniques for analyzing single-cell RNA sequencing data. Disp [36] and VST [31]are widely used gene filtering methods. Disp, introduced by Seurat, identifies genes with the largest variation after controlling for mean expression variability by z-standardizing dispersion measures within expression bins. VST refines this approach by fitting a loess curve to the log(variance) vs. log(mean) relationship.

In terms of gene selection methods, SAIC [64] uses an iterative k-means clustering method to thoroughly search for the best feature genes. FEAST [38] uses the F statistic to test feature significance and summarize the variance differences between and within groups, similar to the Fisher score. We also selected the ensemble learning-based method, CellBRF [65], which uses a random forest guided by predicted cell labels to identify the most important genes for distinguishing cell types. Finally, we considered the graph-based method, HRG [39], which finds informative genes by optimizing expression patterns in a similarity network between cells, ensuring that these genes exhibit regional expression patterns. Each method provides a unique approach to gene selection.

### G. Benchmarks Data

We selected 36 benchmark datasets with cell numbers ranging from 90 to 265,767, covering various tissues of humans and mice, including pancreas, brain, intestine, spleen, liver, bone marrow, retina, etc. These datasets also involve a variety of diseases and health conditions, such as human islet cells, mouse cerebral cortex, human cervical cancer, and mouse motor cortex. The number of genes in these datasets ranges from 14,717 to 59,357, and the number of cell types ranges from 3 to 20, covering a wide range of biodiversity to measure the performance of CellScope. The detailed information of all datasets are listed in **Supplementary Table S1**.

## Data availability

Publicly available datasets used in this study can be accessed from the National Center for Biotechnology Information Gene Expression Omnibus(GSE36552, GSE83139, GSE67835, GSE59739, GSE81608, GSE85241, GSE65525, GSE60361, GSE132042, GSE108097, GSE103322, GSE84133, GSE178101, GSE228590, GSE160189, GSE243413 and GSE178101), the European Nucleotide Archive (E-MTAB-3321, E-MTAB-13382, E-MTAB-12795 and E-MTAB-10187), the Database of Genotypes and Phenotypes(PHS000833, PHS000424V9P2), and the BRAIN Initiative Cell Census Network (RRID:SCR_015820) and are available for download from the Neuroscience Multi-omics Archive (RRID:SCR_016152). The specific download link can be found in **Supplementary Table S2**.

## Code availability

CellScope is implemented in Python and available on GitHub at https://github.com/zhigang-yao/CellScope. Detailed tutorials, code instructions and notebooks to reproduce the results of this study are available at https://cellscope.readthedocs.io/en/latest/.

## Acknowledgements

The whole work has been supported by the Singapore Ministry of Education Tier 2 grant (A-0008520-00-00, A-8001562-00-00) and the Tier 1 grant (A8000987-00-00 and A-8002931-00-00) at the National University of Singapore.

## Author contributions

L.B. contributed to literature collection, project planning and designing, framework development, code implementation, figure creation, and paper writing. R.L. conducted literature collection, participated in framework development and experimental code debugging, and contributed to figure creation, and paper writing. T.N. implemented the Python version of the original code, expanded the algorithm’s functionality, and contributed to experimental code debugging, figure creation, and paper writing. G.Y. reviewed the article, provided suggestions for experimental design and article writing. B. M. reviewed the article, helped revise the article. Z.Y. and L.J. developed the idea, supervised the project and wrote the manuscript. All co-authors read and approved the paper.

