## Supplementary Material for "CellScope: High-Performance Cell Atlas Workflow with Tree-Structured Representation"

### Appendix

**References** **28**

### 1 Supplementary Note

#### 1.1 Clustering Validation Indices

We employ five distinct metrics to evaluate the performance of clustering: Adjusted Rand Index (ARI), Accuracy (ACC), Normalized Mutual Information (NMI), Jaccard Index (JI), and F1 Score.

##### 1.1.1 ARI

The Rand Index (RI) [1] quantifies the agreement between a given clustering and the ground truth, calculated as:

$$\mathrm{RI} = \frac{a+b}{a+b+c+d} = \frac{a+b}{\binom{N}{2}} \tag{1}$$

where $a$ is the number of pairs in the same true cell type and clustered together, $b$ is the number of pairs in different true cell types and not clustered together, $c$ is the number of pairs in the same cell type but not clustered together, $d$ is the number of pairs in different cell types but clustered together, and $\binom{N}{2}$ is the total number of possible pairs from $N$ cells. The ARI is a chance-corrected version of RI, ranging from -1 to 1, with 0 indicating random subtyping.

##### 1.1.2 ACC

ACC [2] measures the proportion of correctly classified cells in the clustering:

$$\mathrm{ACC} = \frac{\sum_{i=1}^{n} \delta(\mathrm{map}(c_i), l_i)}{n} \tag{2}$$

where $n$ is the total number of cells, $c_i$ is the cluster assignment of the $i$-th cell, $l_i$ is the true label, $\delta$ is the Kronecker delta function, and map() is a mapping function that permutes cluster labels to match ground truth labels.

##### 1.1.3 NMI

NMI is a normalized version of Mutual Information (MI):

$$\mathrm{NMI} = \frac{1}{2} \times \frac{I(X;Y)}{H(X)+H(Y)} \tag{3}$$

where $X$ is the true labeling, $Y$ is the clustering partition, $I(X;Y)$ is the mutual information between $X$ and $Y$, and $H(X)$ and $H(Y)$ are the entropies of $X$ and $Y$, respectively. NMI ranges from 0 to 1, with 1 indicating perfect matching between cell types and clusters.

### 1.1.4 JI

Also known as Intersection over Union, JI [3] is calculated as:

$$\mathrm{JI} = \frac{a}{a+b+c} \tag{4}$$

where $a$, $b$, and $c$ are defined as in the RI calculation. These metrics provide a comprehensive evaluation of clustering performance, each capturing different aspects of the agreement between algorithm-generated clusters and true cell types.

##### 1.1.5 F1 Score

The F1 score, a widely used metric in clustering evaluation, is defined as the harmonic mean of precision and recall [4]. It is calculated as:

$$\text{F1 score} = 2 \cdot \frac{\text{Precision} \cdot \text{Recall}}{\text{Precision} + \text{Recall}} \tag{5}$$

where:

- **Precision**: The proportion of correctly predicted positive instances out of all predicted positive instances.
- **Recall**: The proportion of correctly predicted positive instances out of all actual positive instances.

##### 1.2 Gene Selection Evaluation Metrics

- **Neighborhood Purity**: Measuring the local homogeneity of cell types around each cell by calculating the proportion of a cell's neighbor cells that share the same cell type, reflecting the purity of local neighborhoods.
- **Variance Ratio**: The ratio of between-cell-type variance to within-cell-type variance, measuring the separation of different cell types.
- **Average Silhouette Width (ASW) [5]**: Measuring the separation of different cell types by comparing the average distance of a cell to others of the same cell type versus the average distance to cells from the closest other cell type.
- **KNN Classification Accuracy**: The accuracy for predicting a cell's type based on the cell type labels of its k nearest neighbor cells, reflects the clustering of similar cells.
- **Cell-type Local Inverse Simpson Index (LISI) [6]**: Characterizing the local diversity of cell types around a cell, calculated by the expected number of draws needed to obtain at least one cell with the same cell type from a cell's neighbors.

##### 1.3 Baseline Methods

###### 1.3.1 Compared Pipelines

We compared CellScope with two existing popular baseline methods, Scanpy [7] and Seurat [8] in this study. Scanpy was implemented from its original source code repository (https://github.com/scverse/Scanpy). Highly variable genes were identified based on specified thresholds for mean expression and dispersion, and clustering was performed using the Leiden algorithm across a range of resolutions. The algorithm parameters were set according to the default parameter settings in the tutorial(https://Scanpy-tutorials.readthedocs.io/en/latest/pbmc3k.html). Then, we calculated the adjusted Rand index (ARI) between the predicted clusters and the true cell type labels for each resolution.

Seurat was implemented from its source code (https://satijalab.org/seurat) with a scale factor of 10,000. We identified 2,000 variable features using the VST selection method as mentioned in Seurat tutorial(https://satijalab.org/seurat/articles/tutorial). Neighbors were identified using the first 10 principal components, and clustering by Louvain algorithm was performed across a range of resolutions. For each clustering resolution, we calculated the ARI between the predicted clusters and the true cell type labels.

###### 1.3.2 Gene Selection Methods

We compared several common gene filtering methods and the latest gene selection methods to identify the most effective techniques for analyzing single-cell RNA sequencing data. Disp [9] and VST [8]are widely used gene filtering methods. Disp, introduced by Seurat, identifies genes with the largest variation after controlling for mean expression variability by z-standardizing dispersion measures within expression bins. VST refines this approach by fitting a loess curve to the log(variance) vs. log(mean) relationship. We also consider HVG methods provided by Scanpy [7], considering the normalized dispersion obtained by scaling with the mean and standard deviation of the dispersions for genes falling into a given bin for the mean expression of genes.

In terms of gene selection methods, FEAST [10] uses the F statistic to test feature significance and summarize the variance differences between and within groups, similar to the Fisher score. We also selected the mixture of multiple highly variable gene selection methods, mixHVG [11], which can combine the advantages captured by every single method. Finally, we considered the graph-based method, HRG [12], which finds informative genes by optimizing expression patterns in a similarity network between cells, ensuring that these genes exhibit regional expression patterns. Each method provides a unique approach to gene selection.

##### 1.4 Differential Expression Analysis Method

According to the article, we define the cell dataset $\mathcal{X} = \{x_i\}_{i=1}^N$, where $\mathcal{X}$ represents a collection of single-cell RNA sequencing raw data. Each vector $x_i$ corresponds to the expression values $[x_i^{(1)}, \dots, x_i^{(D)}]$ of the $i$-th cell across $D$ genes. We denote these two classes of cells as $C_1$ and $C_2$, comprising $N_1$ and $N_2$ cells, respectively. Next, we will introduce four methods to identify differentially expressed genes between these two cell types.

##### 1.4.1 Fold Change Method

The fold change method calculates the ratio of average gene expression between $C_1$ and $C_2$, highlighting genes with significant proportional differences:

$$FC_j = \left| \log_2 \left( 1 + \frac{\frac{1}{N_1} \sum_{x_i \in C_1} x_i^{(j)}}{\frac{1}{N_2} \sum_{x_i \in C_2} x_i^{(j)}} \right) \right|, \ \ j = 1, \cdots, D$$

Furthermore, we select the gene corresponding to the highest $FC$ as the differentially expressed gene between the two classes $C_1$ and $C_2$.

##### 1.4.2 Gene Expression Percentage

The percentage difference method evaluates the relative frequency of non-zero expression values within each population, highlighting genes with distinct distribution patterns. Specifically, for each gene, the metric is calculated as:

$$pct_j = \left| \frac{1}{N_1} \sum_{x_i \in C_1} sgn(x_i^{(j)}) - \frac{1}{N_2} \sum_{x_i \in C_2} sgn(x_i^{(j)}) \right|,$$

where $sgn(\cdot)$ is the sign function defined as

$$\text{sgn}(x) = \begin{cases} 1, & \text{if } x \neq 0 \\ 0, & \text{if } x = 0 \end{cases}$$

Furthermore, we select the gene corresponding to the highest *pct* as the differentially expressed gene between the two classes $C_1$ and $C_2$.

Following these analyses, we employ a rigorous sorting and filtering mechanism to identify genes with the highest differential scores, thereby offering robust candidates as marker genes. This method not only enhances our understanding of cellular functions but also provides valuable insights into disease mechanisms. These findings are crucial for advancing research in precision medicine and therapeutic development.

##### 1.4.3 Wilcoxon Rank-Sum Test

The Wilcoxon rank-sum test, also known as the Mann-Whitney U test, is a non-parametric test used to compare whether the medians of two independent samples are the same. Below are the basic steps and formulas for the Wilcoxon rank-sum test:

Let $C_1 = \left\{ x_{i_1}, \cdots, x_{i_{N_1}} \right\}$ and $C_2 = \left\{ x_{j_1}, \cdots, x_{j_{N_2}} \right\}$. For gene $k$, let $C_{1k} = \left\{ x_{i_1}^{(k)}, \cdots, x_{i_{N_1}}^{(k)} \right\}$ be the expression values of gene $k$ in cell type $C_1$, and $C_{2k} = \left\{ x_{j_1}^{(k)}, \cdots, x_{j_{N_2}}^{(k)} \right\}$ be the sample of expression values in cell type $C_2$. We then combine the two samples $C_{1k}$ and $C_{2k}$ into a single sample $X_k = \{C_{1k}, C_{2k}\}$. Rank the combined data from smallest to largest, and assign ranks to the combined sample.

Let $R_1$ be the sum of the ranks for the sample $C_{1k}$ and $R_2$ be the sum of the ranks for the sample $C_{2k}$. Calculate the test statistic $U$ for both groups as:

$$U_1 = R_1 - \frac{N_1(N_1 + 1)}{2}$$

$$U_2 = R_2 - \frac{N_2(N_2 + 1)}{2}$$

Let $U = \min\{U_1, U_2\}$. Compare the test statistic $U$ to the critical value from the Mann-Whitney U distribution table, or calculate the p-value.

After computing the p-values for each gene, select the top $K$ genes with the smallest p-values as the differentially expressed genes.

##### 1.4.4 t-Test

The t-test is a parametric test used to compare whether the means of two independent samples are the same. To identify differentially expressed genes between two cell types $C_1$ and $C_2$, we start by defining the sets $C_1 = \left\{x_{i_1}, \cdots, x_{i_{N_1}}\right\}$ and $C_2 = \left\{x_{j_1}, \cdots, x_{j_{N_2}}\right\}$. For a specific gene $k$, the expression values in cell type $C_1$ are $C_{1k} = \left\{x_{i_1}^{(k)}, \cdots, x_{i_{N_1}}^{(k)}\right\}$ and in cell type $C_2$ are $C_{2k} = \left\{x_{j_1}^{(k)}, \cdots, x_{j_{N_2}}^{(k)}\right\}$.

First, we calculate the mean expression levels for gene $k$ in both cell types, denoted as $\bar{X}_{1k}$ and $\bar{X}_{2k}$. Next, we compute the variances for gene $k$ in both cell types, denoted as $s_{1k}^2$ and $s_{2k}^2$. Using these variances, we calculate the pooled standard deviation $s_p$, which accounts for the variances in both cell types. The t-statistic for gene $k$ is then calculated using the formula $t_k = \frac{\bar{X}_{1k} - \bar{X}_{2k}}{s_p\sqrt{\frac{1}{N_1} + \frac{1}{N_2}}}$.

The t-statistic allows us to determine the p-value from the t-distribution with $N_1 + N_2 - 2$ degrees of freedom, indicating whether the difference in means is statistically significant. When testing multiple genes, we adjust the p-values to account for multiple comparisons, using methods such as the Bonferroni correction or the Benjamini-Hochberg procedure. Finally, we select the genes with adjusted p-values below a chosen significance threshold (e.g., 0.05) as differentially expressed genes.

### 2 Supplementary Table

#### 2.1 Data Information

Table S1: Information of 36 Benchmark scRNA-seq Datasets

| Data Codes | Data Name | Tissue | Year | #Cells | #Genes | #Cell types |
|---|---|---|---|---|---|---|
| GSE36552 | Yan | Human Embryos | 2013 | 90 | 20214 | 6 |
| E-MTAB-3321 | Goolam | Mouse Embryos | 2016 | 124 | 41428 | 5 |
| SRP041736 | Pollen | Human Brain | 2014 | 249 | 14805 | 11 |
| GSE83139 | Wang | Human Pancreatic | 2016 | 457 | 19950 | 7 |
| GSE67835 | Darmanis | Human Brain | 2015 | 466 | 22088 | 9 |
| GSE59739 | Usoskin | Mouse Lumbar | 2015 | 622 | 25334 | 4 |
| GSE81608 | Xin | Human Islet | 2016 | 1600 | 39851 | 8 |
| GSE132042-Intestine | Keller(I) | Mouse Intestine | 2019 | 1887 | 17985 | 5 |
| GSE85241 | Muraro | Human Pancreas | 2016 | 2126 | 19127 | 10 |
| PHS000424V9P2-Brain | NHGRI | Human Brain | 2022 | 2642 | 14717 | 8 |
| GSE65525 | Klein | Mouse Embryos | 2015 | 2717 | 24175 | 4 |
| GSE60361 | Zeisel | Mouse Cerebral Cortex | 2015 | 3005 | 19972 | 7 |
| PHS000833 | Lake | Human Brain | 2018 | 3042 | 25051 | 16 |
| GSE132042-Pancreas | Keller(P) | Mouse Pancreas | 2019 | 3384 | 21069 | 9 |
| GSE108097-Brain | Han(B) | Mouse Brain | 2018 | 4038 | 16906 | 15 |
| SCR015820-MidBrain | Siletti-1 | Human Brain | 2023 | 4714 | 59357 | 11 |
| GSE103322 | Tirosh | Human oral cavity | 2017 | 5902 | 23686 | 10 |
| SCR015820-LNC | Siletti-2 | Human Brain | 2023 | 6877 | 59357 | 10 |
| GSE84133-Human | Baron(H) | Human Pancreas | 2016 | 8569 | 20125 | 14 |
| SCR015820-Pons | Siletti-3 | Human Brain | 2023 | 23349 | 59236 | 13 |
| SCR015820-THM | Siletti-4 | Human Brain | 2023 | 27111 | 59236 | 11 |
| SCR015820-HiB | Siletti-5 | Human Brain | 2023 | 28724 | 59236 | 12 |
| SCR015820-Pn | Siletti-6 | Human Brain | 2023 | 47416 | 59236 | 13 |
| GSE178101 | Ulrich(H) | Human Fallopian Tubes | 2021 | 59738 | 36960 | 12 |
| E-MTAB-13382 | Li | Human Trophoblast Organoids | 2024 | 67996 | 36398 | 3 |
| — | Li(H) | Human Retina | 2023 | 72788 | 36398 | 13 |
| GSE228590-LA | Qiu(L) | Mouse Embryonic | 2024 | 76732 | 45854 | 6 |
| GSE178101 | Ulrich | Human Fallopian Tubes | 2021 | 77536 | 36960 | 12 |
| GSE228590-EO | Qiu(E) | Mouse Embryonic | 2024 | 93695 | 45854 | 11 |
| GSE160189 | Ayhan | Hippocampal | 2021 | 129905 | 17132 | 8 |
| GSM5027160 etc | Posner | Human Brain | 2022 | 130908 | 25232 | 16 |
| GSE243413 | Li(M) | Mouse Retina | 2024 | 147523 | 32034 | 13 |
| E-MTAB-12795 | Hoo | Human Placenta | 2024 | 158978 | 36398 | 7 |
| E-MTAB-10187 | Yu | Human Multi-Endodermal | 2021 | 155232 | 30867 | 20 |
| SCR016152-2 | Yao | Mouse Motor Cortex | 2021 | 159738 | 30511 | 11 |
| — | Li(HA) | Human Retina | 2023 | 265767 | 36398 | 18 |

##### 2.2 Data Source

Table S2: Download Links of 36 Benchmark scRNA-seq Datasets

| Data Name | Access Link |
|---|---|
| Yan | https://www.ncbi.nlm.nih.gov/geo/query/acc.cgi?acc=GSE36552 |
| Goolam | https://www.ebi.ac.uk/biostudies/arrayexpress/studies/E-MTAB-3321 |
| Pollen | https://www.ncbi.nlm.nih.gov/geo/query/acc.cgi?acc=GSM1832359 |
| Wang | https://www.ncbi.nlm.nih.gov/geo/query/acc.cgi?acc=GSE83139 |
| Darmanis | https://www.ncbi.nlm.nih.gov/geo/query/acc.cgi?acc=GSE67835 |
| Usoskin | https://www.ncbi.nlm.nih.gov/geo/query/acc.cgi?acc=GSE59739 |
| Xin | https://www.ncbi.nlm.nih.gov/geo/query/acc.cgi?acc=GSE81608 |
| Keller(I) | https://cellxgene.cziscience.com/collections/0b9d8a04-bb9d-44da-aa27-705bb65b54eb |
| Muraro | https://cellxgene.cziscience.com/collections/6e8c5415-302c-492a-a5f9-f29c57ff18fb |
| NHGRI | gtexportal.org/home/aboutAdultGtex |
| Klein | https://www.ncbi.nlm.nih.gov/geo/query/acc.cgi?acc=GSE65525 |
| Zeisel | https://www.ncbi.nlm.nih.gov/geo/query/acc.cgi?acc=GSM1474572 |
| Lake | https://www.ncbi.nlm.nih.gov/projects/gap/cgi-bin/study.cgi?study_id=phs000833.v3.p1 |
| Keller(P) | https://cellxgene.cziscience.com/collections/0b9d8a04-bb9d-44da-aa27-705bb65b54eb |
| Han(B) | https://www.ncbi.xyz/geo/query/acc.cgi?acc=GSM2906452 |
| Siletti-1 | https://cellxgene.cziscience.com/collections/283d65eb-dd53-496d-adb7-7570c7caa443 |
| Tirosh | https://www.ncbi.nlm.nih.gov/geo/query/acc.cgi?acc=GSE103322 |
| Siletti-2 | https://cellxgene.cziscience.com/collections/283d65eb-dd53-496d-adb7-7570c7caa443 |
| Baron(H) | https://www.ncbi.nlm.nih.gov/geo/query/acc.cgi?acc=GSE84133 |
| Siletti-3 | https://cellxgene.cziscience.com/collections/283d65eb-dd53-496d-adb7-7570c7caa443 |
| Siletti-4 | https://cellxgene.cziscience.com/collections/c26ca66a-63ea-4059-a24e-0e0be0a2a173 |
| Siletti-5 | https://cellxgene.cziscience.com/collections/283d65eb-dd53-496d-adb7-7570c7caa443 |
| Siletti-6 | https://cellxgene.cziscience.com/collections/283d65eb-dd53-496d-adb7-7570c7caa443 |
| Ulrich(H) | https://cellxgene.cziscience.com/collections/fc77d2ae-247d-44d7-aa24-3f4859254c2c |
| Li | https://cellxgene.cziscience.com/collections/257ce7fb-ab27-4772-8a06-9fe8385816b2 |
| Li(H) | https://cellxgene.cziscience.com/collections/4c6eaf5c-6d57-4c76-b1e9-60df8c655f1e |
| Qiu(L) | https://cellxgene.cziscience.com/collections/45d5d2c3-bc28-4814-aed6-0bb6f0e11c82 |
| Ulrich | https://cellxgene.cziscience.com/collections/fc77d2ae-247d-44d7-aa24-3f4859254c2c |
| Qiu(E) | https://cellxgene.cziscience.com/collections/45d5d2c3-bc28-4814-aed6-0bb6f0e11c82 |
| Ayhan | https://cellxgene.cziscience.com/collections/f17b9205-f61f-4a0f-a65a-73ba91c50ade |
| Posner | https://cellxgene.cziscience.com/collections/e9eec7f5-8519-42f6-99b4-6dbd9cc5ef03 |
| Li(M) | https://cellxgene.cziscience.com/collections/a0c84e3f-a5ca-4481-b3a5-ccfda0a81ecc |
| Hoo | https://cellxgene.cziscience.com/collections/5f80428b-222d-450b-a7de-a408186ceb86 |
| Yu | https://cellxgene.cziscience.com/collections/dfc09a93-bce0-4c77-893d-e153d1b7f9fa |
| Yao | https://cellxgene.cziscience.com/collections/ae1420fe-6630-46ed-8b3d-cc6056a66467 |
| Li(HA) | https://cellxgene.cziscience.com/collections/4c6eaf5c-6d57-4c76-b1e9-60df8c655f1e |

#### 2.3 Clustering Performance Comparison

Table S3: Clustering Performance on 36 Benchmark scRNA-seq Datasets Using CellScope, Seurat, and Scanpy, Measured by ARI, NMI, ACC, and JI

| Data Name | ARI | | | NMI | | | ACC | | | JI | | |
|---|---|---|---|---|---|---|---|---|---|---|---|---|
| | CellScope | Seurat | Scanpy | CellScope | Seurat | Scanpy | CellScope | Seurat | Scanpy | CellScope | Seurat | Scanpy |
| Yan | 0.97 | 0.86 | 0.85 | 0.97 | 0.89 | 0.88 | 0.93 | 0.82 | 0.86 | 0.88 | 0.70 | 0.75 |
| Goolam | 0.91 | 0.86 | 0.63 | 0.91 | 0.82 | 0.69 | 0.82 | 0.82 | 0.77 | 0.70 | 0.70 | 0.63 |
| Pollen | 0.90 | 0.89 | 0.86 | 0.94 | 0.91 | 0.91 | 0.91 | 0.90 | 0.86 | 0.83 | 0.82 | 0.75 |
| Wang | 0.86 | 0.82 | 0.62 | 0.85 | 0.78 | 0.65 | 0.91 | 0.90 | 0.84 | 0.84 | 0.81 | 0.72 |
| Darmanis | 0.76 | 0.69 | 0.46 | 0.77 | 0.75 | 0.60 | 0.85 | 0.75 | 0.62 | 0.74 | 0.60 | 0.45 |
| Usoskin | 0.97 | 0.79 | 0.74 | 0.94 | 0.77 | 0.80 | 0.98 | 0.86 | 0.82 | 0.97 | 0.76 | 0.69 |
| Xin | 0.92 | 0.94 | 0.95 | 0.83 | 0.83 | 0.87 | 0.92 | 0.92 | 0.93 | 0.86 | 0.85 | 0.87 |
| Keller(I) | 0.80 | 0.70 | 0.78 | 0.69 | 0.65 | 0.69 | 0.84 | 0.80 | 0.84 | 0.73 | 0.66 | 0.73 |
| Muraro | 0.92 | 0.92 | 0.49 | 0.88 | 0.88 | 0.69 | 0.95 | 0.95 | 0.63 | 0.90 | 0.91 | 0.46 |
| NHGRI | 0.95 | 0.82 | 0.77 | 0.92 | 0.78 | 0.76 | 0.97 | 0.82 | 0.82 | 0.94 | 0.69 | 0.69 |
| Klein | 0.97 | 0.90 | 0.91 | 0.95 | 0.90 | 0.92 | 0.98 | 0.92 | 0.94 | 0.97 | 0.86 | 0.88 |
| Zeisel | 0.80 | 0.75 | 0.69 | 0.77 | 0.72 | 0.71 | 0.87 | 0.87 | 0.83 | 0.76 | 0.77 | 0.71 |
| Lake | 0.82 | 0.76 | 0.64 | 0.78 | 0.79 | 0.70 | 0.79 | 0.77 | 0.65 | 0.65 | 0.63 | 0.48 |
| Keller(P) | 0.88 | 0.79 | 0.09 | 0.85 | 0.78 | 0.25 | 0.92 | 0.84 | 0.36 | 0.86 | 0.73 | 0.22 |
| Han(B) | 0.89 | 0.16 | 0.36 | 0.72 | 0.25 | 0.62 | 0.91 | 0.41 | 0.60 | 0.83 | 0.26 | 0.43 |
| Siletti-1 | 0.93 | 0.93 | 0.77 | 0.93 | 0.93 | 0.85 | 0.95 | 0.95 | 0.85 | 0.90 | 0.90 | 0.74 |
| Tirosh | 0.90 | 0.50 | 0.56 | 0.87 | 0.74 | 0.76 | 0.92 | 0.62 | 0.63 | 0.86 | 0.45 | 0.46 |
| Siletti-2 | 0.99 | 0.87 | 0.87 | 0.97 | 0.88 | 0.88 | 0.98 | 0.86 | 0.87 | 0.97 | 0.76 | 0.77 |
| Baron(H) | 0.92 | 0.95 | 0.91 | 0.89 | 0.92 | 0.88 | 0.94 | 0.94 | 0.91 | 0.88 | 0.89 | 0.83 |
| Siletti-3 | 0.84 | 0.51 | 0.55 | 0.89 | 0.76 | 0.79 | 0.92 | 0.66 | 0.68 | 0.86 | 0.49 | 0.52 |
| Siletti-4 | 0.99 | 0.39 | 0.39 | 0.97 | 0.70 | 0.70 | 0.98 | 0.54 | 0.54 | 0.96 | 0.37 | 0.37 |
| Siletti-5 | 0.88 | 0.16 | 0.20 | 0.84 | 0.50 | 0.52 | 0.95 | 0.47 | 0.51 | 0.90 | 0.31 | 0.34 |
| Siletti-6 | 0.90 | 0.35 | 0.62 | 0.88 | 0.66 | 0.78 | 0.94 | 0.46 | 0.74 | 0.88 | 0.30 | 0.59 |
| Ulrich(H) | 0.80 | 0.63 | 0.77 | 0.86 | 0.80 | 0.85 | 0.85 | 0.74 | 0.84 | 0.73 | 0.58 | 0.72 |
| Li | 0.86 | 0.44 | 0.37 | 0.68 | 0.49 | 0.56 | 0.91 | 0.65 | 0.59 | 0.83 | 0.48 | 0.42 |
| Li(H) | 0.95 | 0.70 | 0.90 | 0.94 | 0.86 | 0.93 | 0.95 | 0.74 | 0.89 | 0.91 | 0.59 | 0.80 |
| Qiu(L) | 0.75 | 0.71 | 0.68 | 0.73 | 0.77 | 0.77 | 0.89 | 0.74 | 0.73 | 0.80 | 0.59 | 0.58 |
| Ulrich | 0.79 | 0.63 | 0.77 | 0.87 | 0.80 | 0.86 | 0.86 | 0.74 | 0.86 | 0.76 | 0.59 | 0.76 |
| Qiu(E) | 0.81 | 0.73 | 0.52 | 0.77 | 0.74 | 0.69 | 0.86 | 0.83 | 0.70 | 0.75 | 0.71 | 0.54 |
| Ayhan | 0.88 | 0.82 | 0.69 | 0.81 | 0.80 | 0.85 | 0.91 | 0.85 | 0.84 | 0.83 | 0.73 | 0.73 |
| Posner | 0.82 | 0.75 | 0.69 | 0.84 | 0.82 | 0.78 | 0.78 | 0.74 | 0.64 | 0.63 | 0.58 | 0.47 |
| Li(M) | 0.80 | 0.67 | 0.79 | 0.87 | 0.84 | 0.92 | 0.82 | 0.73 | 0.80 | 0.70 | 0.57 | 0.67 |
| Hoo | 0.96 | 0.94 | 0.83 | 0.95 | 0.95 | 0.89 | 0.98 | 0.96 | 0.84 | 0.96 | 0.93 | 0.73 |
| Yu | 0.86 | 0.37 | 0.72 | 0.80 | 0.71 | 0.78 | 0.83 | 0.59 | 0.81 | 0.70 | 0.41 | 0.68 |
| Yao | 0.83 | 0.35 | 0.39 | 0.81 | 0.74 | 0.76 | 0.81 | 0.54 | 0.60 | 0.68 | 0.37 | 0.43 |
| Li(HA) | 0.88 | 0.57 | 0.80 | 0.88 | 0.79 | 0.85 | 0.82 | 0.64 | 0.77 | 0.69 | 0.47 | 0.62 |
| Average | 0.88 | 0.68 | 0.66 | 0.86 | 0.77 | 0.76 | 0.90 | 0.76 | 0.75 | 0.82 | 0.63 | 0.62 |

#### 2.4 Runtime Comparison

Table S4: CPU Runtime on 36 Benchmark scRNA-seq Datasets Using CellScope, Scanpy, and Seurat

| Data Name | #Cells | #Classes | CellScope (s) | Scanpy (s) | Seurat (s) | Scanpy/CellScope | Seurat/CellScope |
|---|---|---|---|---|---|---|---|
| Yan | 90 | 6 | 0.93 | 0.34 | 10.83 | 0.37 | 11.65 |
| Goolam | 124 | 5 | 1.45 | 0.46 | 15.25 | 0.32 | 10.52 |
| Pollen | 249 | 11 | 1.73 | 0.79 | 18.68 | 0.46 | 10.80 |
| Wang | 457 | 7 | 1.62 | 1.98 | 23.20 | 1.22 | 14.32 |
| Darmanis | 466 | 9 | 1.25 | 1.92 | 15.16 | 1.54 | 12.13 |
| Usoskin | 622 | 4 | 3.44 | 2.20 | 17.79 | 0.64 | 5.17 |
| Xin | 1600 | 8 | 4.24 | 9.57 | 44.64 | 2.26 | 10.53 |
| Keller(I) | 1887 | 5 | 3.19 | 8.92 | 36.94 | 2.80 | 11.58 |
| Muraro | 2126 | 10 | 2.68 | 7.59 | 47.94 | 2.83 | 17.89 |
| NHGRI | 2642 | 8 | 2.67 | 10.71 | 52.37 | 4.01 | 19.61 |
| Klein | 2717 | 4 | 4.89 | 13.18 | 68.50 | 2.70 | 14.01 |
| Zeisel | 3005 | 7 | 4.23 | 12.14 | 59.50 | 2.87 | 14.07 |
| Lake | 3042 | 16 | 4.48 | 16.96 | 83.55 | 3.79 | 18.65 |
| Keller(P) | 3384 | 9 | 4.25 | 15.81 | 86.98 | 3.72 | 20.47 |
| Han(B) | 4038 | 15 | 3.86 | 23.80 | 205.15 | 6.17 | 53.15 |
| Siletti-1 | 4714 | 11 | 7.56 | 25.64 | 106.29 | 3.39 | 14.06 |
| Tirosh | 5902 | 10 | 6.31 | 24.42 | 110.29 | 3.87 | 17.48 |
| Siletti-2 | 6877 | 10 | 9.92 | 45.48 | 132.60 | 4.58 | 13.37 |
| Baron(H) | 8569 | 14 | 8.11 | 40.80 | 129.84 | 5.03 | 16.01 |
| Siletti-3 | 23349 | 13 | 61.76 | 180.07 | 350.66 | 2.92 | 5.68 |
| Siletti-4 | 27111 | 11 | 88.09 | 181.37 | 321.97 | 2.06 | 3.66 |
| Siletti-5 | 28724 | 12 | 90.09 | 213.37 | 301.97 | 2.37 | 3.35 |
| Siletti-6 | 47416 | 13 | 158.78 | 467.33 | 740.04 | 2.94 | 4.66 |
| Ulrich(H) | 59738 | 12 | 276.53 | 405.73 | 722.65 | 1.47 | 2.61 |
| Li | 67996 | 3 | 220.93 | 454.89 | 807.52 | 2.06 | 3.66 |
| Li(H) | 72788 | 13 | 312.74 | 596.29 | 849.21 | 1.91 | 2.72 |
| Qiu(L) | 76732 | 6 | 329.69 | 628.60 | 895.22 | 1.91 | 2.72 |
| Ulrich | 77536 | 12 | 508.28 | 649.82 | 1055.31 | 1.28 | 2.08 |
| Qiu(E) | 93695 | 11 | 863.78 | 1397.73 | 1716.79 | 1.62 | 1.99 |
| Ayhan | 129905 | 8 | 892.31 | 2252.85 | 1711.64 | 2.52 | 1.92 |
| Posner | 130908 | 16 | 822.35 | 1550.37 | 1904.30 | 1.89 | 2.32 |
| Li(M) | 147523 | 13 | 916.54 | 1470.60 | 2192.31 | 1.60 | 2.39 |
| Hoo | 158978 | 7 | 1117.50 | 2423.69 | 2334.78 | 2.17 | 2.09 |
| Yu | 155232 | 20 | 1091.17 | 2366.58 | 2279.77 | 2.17 | 2.09 |
| Yao | 159738 | 11 | 1377.25 | 2353.77 | 17659.06 | 1.71 | 12.82 |
| Li(HA) | 265767 | 18 | 1423.72 | 4112.64 | 21154.79 | 2.89 | 14.86 |

##### 2.5 Gene Selection Algorithms Comparison

Table S5: Gene Selection Algorithm Performance on 36 Benchmark scRNA-seq Datasets, Measured by Variance Ratio

| Data Name | Variance Ratio | | | | | | |
|---|---|---|---|---|---|---|---|
| | CellScope | Scanpy | Disp | VST | Mixhvg | FEAST | HRG |
| Yan | 93.15 | 28.31 | 48.12 | 37.41 | 51.61 | 23.92 | 26.55 |
| Goolam | 30.73 | 10.27 | 17.66 | 17.46 | 17.55 | 8.04 | 11.67 |
| Pollen | 30.38 | 8.82 | 10.63 | 11.40 | 14.88 | 49.38 | 46.21 |
| Wang | 40.94 | 6.88 | 12.82 | 13.25 | 13.80 | 64.75 | 54.19 |
| Darmanis | 29.21 | 7.59 | 11.42 | 13.19 | 12.39 | 7.14 | 38.06 |
| Usoskin | 46.48 | 10.20 | 13.50 | 13.10 | 16.92 | 8.46 | 8.25 |
| Xin | 24.42 | 7.13 | 14.25 | 16.93 | 13.88 | 12.07 | 0.98 |
| Keller(I) | 95.37 | 13.46 | 2.92 | 13.50 | 26.23 | 21.07 | 48.00 |
| Muraro | 159.74 | 133.93 | 119.90 | 141.55 | 141.07 | 41.94 | 60.93 |
| NHGRI | 816.59 | 324.23 | 392.86 | 263.67 | 453.54 | 219.95 | 253.09 |
| Klein | 407.98 | 81.33 | 182.34 | 233.52 | 202.18 | 140.50 | 193.76 |
| Zeisel | 447.95 | 131.78 | 218.83 | 238.62 | 211.27 | 204.75 | 157.20 |
| Lake | 110.45 | 15.52 | 34.71 | 30.45 | 41.95 | 100.78 | 43.81 |
| Keller(P) | 222.08 | 16.99 | 8.51 | 5.43 | 35.40 | 41.09 | 85.05 |
| Han(B) | 167.55 | 68.93 | 154.19 | 130.17 | 129.72 | | 113.81 |
| Siletti-1 | 710.29 | 208.19 | 151.45 | 159.62 | 160.61 | | 141.73 |
| Tirosh | 400.43 | 97.55 | 128.15 | 199.72 | 124.24 | | 113.91 |
| Siletti-2 | 728.92 | 192.71 | 534.28 | 387.69 | 486.35 | | 302.34 |
| Baron(H) | 665.53 | 245.72 | 326.28 | 330.44 | 239.68 | | 156.82 |
| Siletti-3 | 3071.74 | 661.67 | 1432.87 | 723.93 | 1186.30 | | 488.81 |
| Siletti-4 | 1697.79 | 612.25 | 1518.73 | 781.53 | 1433.72 | | 547.89 |
| Siletti-5 | 1631.50 | 409.94 | 1099.54 | 726.61 | 1061.26 | | 367.47 |
| Siletti-6 | 4145.32 | 892.88 | 2192.17 | 1447.48 | 2000.17 | | 764.46 |
| Ulrich(H) | 5132.45 | 2746.86 | 3034.92 | 3479.10 | 2551.72 | | |
| Li | 4707.25 | 3117.20 | 3444.75 | 3672.15 | 3011.56 | | |
| Li(H) | 2031.31 | 884.31 | 1894.66 | 1810.87 | 1011.76 | | |
| Qiu(L) | 4027.09 | 1762.96 | 3236.59 | 3085.49 | 1962.67 | | |
| Ulrich | 6685.84 | 3790.12 | 4112.69 | 4826.60 | 3423.98 | | |
| Qiu(E) | 2937.51 | 1477.16 | 2163.92 | 1622.38 | 1368.50 | | |
| Ayhan | 4678.00 | 2221.91 | 4046.85 | 3097.75 | 3067.61 | | |
| Posner | 12474.03 | 4254.84 | 7456.80 | 7060.53 | 5571.82 | | |
| Li(M) | 5770.47 | 4585.23 | 7344.53 | 6559.96 | 3449.99 | | |
| Hoo | 19197.59 | 14164.72 | 16221.75 | 14657.54 | 15273.06 | | |
| Yu | 4448.96 | 3056.37 | 3326.15 | 3298.42 | 2745.47 | | |
| Yao | 19499.83 | 6669.75 | 11457.15 | 8759.38 | 11186.54 | | |
| Li(HA) | 12706.09 | 5429.80 | 8868.99 | 5737.67 | 6549.07 | | |
| Average | 3335.30 | 1620.77 | 2367.66 | 2044.57 | 1923.57 | 67.42 | 175.00 |

Table S6: Gene Selection Algorithm Performance on 36 Benchmark scRNA-seq Datasets, Measured by Average Silhouette Width

| Data Name | Average Silhouette Width | | | | | | |
|---|---|---|---|---|---|---|---|
| | CellScope | Scanpy | Disp | VST | Mixhvg | FEAST | HRG |
| Yan | 0.36 | 0.16 | 0.28 | 0.22 | 0.27 | 0.13 | 0.16 |
| Goolam | 0.25 | 0.14 | 0.24 | 0.19 | 0.19 | 0.12 | 0.16 |
| Pollen | 0.27 | 0.10 | 0.13 | 0.14 | 0.18 | 0.35 | 0.36 |
| Wang | 0.09 | 0.03 | 0.05 | 0.06 | 0.05 | 0.13 | 0.08 |
| Darmanis | 0.11 | 0.02 | 0.02 | 0.05 | 0.03 | 0.01 | 0.08 |
| Usoskin | 0.16 | 0.01 | 0.01 | 0.02 | 0.02 | 0.00 | -0.02 |
| Xin | 0.07 | 0.01 | 0.01 | 0.06 | -0.01 | 0.02 | -0.06 |
| Keller(I) | 0.06 | 0.02 | -0.18 | 0.01 | -0.02 | 0.00 | -0.02 |
| Muraro | 0.16 | 0.16 | 0.14 | 0.18 | 0.15 | 0.04 | 0.06 |
| NHGRI | 0.37 | 0.21 | 0.23 | 0.19 | 0.25 | 0.16 | 0.17 |
| Klein | 0.14 | 0.02 | 0.06 | 0.10 | 0.07 | 0.02 | 0.04 |
| Zeisel | 0.15 | 0.06 | 0.12 | 0.15 | 0.11 | 0.09 | 0.09 |
| Lake | 0.07 | -0.01 | 0.01 | 0.01 | 0.01 | 0.05 | 0.01 |
| Keller(P) | 0.09 | -0.04 | -0.16 | -0.06 | -0.10 | 0.01 | 0.02 |
| Han(B) | 0.09 | -0.11 | 0.13 | 0.17 | 0.10 | | 0.08 |
| Siletti-1 | 0.33 | 0.22 | 0.18 | 0.19 | 0.18 | | 0.16 |
| Tirosh | 0.13 | 0.07 | 0.09 | 0.14 | 0.09 | | 0.07 |
| Siletti-2 | 0.29 | 0.14 | 0.28 | 0.31 | 0.24 | | 0.21 |
| Baron(H) | 0.20 | 0.10 | 0.15 | 0.14 | 0.11 | | 0.08 |
| Siletti-3 | 0.36 | 0.26 | 0.32 | 0.24 | 0.24 | | 0.14 |
| Siletti-4 | 0.29 | 0.18 | 0.31 | 0.19 | 0.29 | | 0.17 |
| Siletti-5 | 0.32 | 0.22 | 0.34 | 0.24 | 0.31 | | 0.19 |
| Siletti-6 | 0.31 | 0.18 | 0.30 | 0.25 | 0.26 | | 0.15 |
| Ulrich(H) | 0.19 | 0.14 | 0.15 | 0.17 | 0.13 | | |
| Li | 0.12 | 0.07 | 0.08 | 0.10 | 0.07 | | |
| Li(H) | 0.10 | 0.04 | 0.10 | 0.09 | 0.05 | | |
| Qiu(L) | 0.13 | 0.06 | 0.11 | 0.10 | 0.06 | | |
| Ulrich | 0.19 | 0.13 | 0.15 | 0.17 | 0.13 | | |
| Qiu(E) | 0.05 | 0.03 | 0.05 | 0.05 | 0.02 | | |
| Ayhan | 0.08 | -0.03 | 0.08 | 0.10 | 0.04 | | |
| Posner | 0.14 | 0.04 | 0.10 | 0.14 | 0.07 | | |
| Li(M) | 0.13 | 0.13 | 0.17 | 0.15 | 0.09 | | |
| Hoo | 0.14 | 0.12 | 0.13 | 0.15 | 0.12 | | |
| Yu | 0.11 | 0.10 | 0.08 | 0.11 | 0.06 | | |
| Yao | 0.29 | 0.18 | 0.23 | 0.21 | 0.21 | | |
| Li(HA) | 0.12 | -0.01 | 0.07 | 0.09 | 0.04 | | |
| Average | 0.18 | 0.09 | 0.13 | 0.13 | 0.11 | 0.08 | 0.10 |

Table S7: Gene Selection Algorithm Performance on 36 Benchmark scRNA-seq Datasets, Measured by KNN Classification Accuracy

| Data Name | KNN Classification Accuracy | | | | | | |
|---|---|---|---|---|---|---|---|
| | CellScope | Scanpy | Disp | VST | Mixhvg | FEAST | HRG |
| Yan | 1.00 | 1.00 | 1.00 | 1.00 | 1.00 | 0.98 | 1.00 |
| Goolam | 0.99 | 0.99 | 1.00 | 0.98 | 1.00 | 0.98 | 1.00 |
| Pollen | 0.95 | 0.84 | 0.86 | 0.86 | 0.90 | 0.96 | 0.93 |
| Wang | 0.97 | 0.93 | 0.94 | 0.96 | 0.95 | 0.97 | 0.95 |
| Darmanis | 0.95 | 0.90 | 0.92 | 0.92 | 0.91 | 0.89 | 0.87 |
| Usoskin | 0.99 | 0.91 | 0.96 | 0.95 | 0.98 | 0.83 | 0.86 |
| Xin | 0.98 | 0.95 | 0.97 | 0.96 | 0.97 | 0.91 | 0.61 |
| Keller(I) | 0.95 | 0.62 | 0.33 | 0.77 | 0.86 | 0.92 | 0.87 |
| Muraro | 0.98 | 0.98 | 0.98 | 0.98 | 0.98 | 0.93 | 0.92 |
| NHGRI | 0.98 | 0.97 | 0.98 | 0.97 | 0.98 | 0.97 | 0.97 |
| Klein | 0.99 | 0.82 | 0.89 | 0.99 | 0.92 | 0.96 | 0.98 |
| Zeisel | 0.96 | 0.89 | 0.95 | 0.95 | 0.95 | 0.91 | 0.95 |
| Lake | 0.96 | 0.76 | 0.87 | 0.92 | 0.88 | 0.95 | 0.90 |
| Keller(P) | 0.97 | 0.56 | 0.59 | 0.65 | 0.48 | 0.70 | 0.95 |
| Han(B) | 0.96 | 0.96 | 0.97 | 0.96 | 0.97 | | 0.97 |
| Siletti-1 | 1.00 | 1.00 | 1.00 | 1.00 | 1.00 | | 1.00 |
| Tirosh | 0.99 | 0.97 | 0.91 | 0.99 | 0.88 | | 0.85 |
| Siletti-2 | 1.00 | 1.00 | 1.00 | 1.00 | 1.00 | | 1.00 |
| Baron(H) | 0.99 | 0.98 | 0.99 | 0.99 | 0.99 | | 0.99 |
| Siletti-3 | 1.00 | 1.00 | 1.00 | 1.00 | 1.00 | | 1.00 |
| Siletti-4 | 1.00 | 0.99 | 1.00 | 1.00 | 1.00 | | 1.00 |
| Siletti-5 | 1.00 | 0.99 | 1.00 | 1.00 | 1.00 | | 1.00 |
| Siletti-6 | 1.00 | 0.99 | 1.00 | 1.00 | 1.00 | | 1.00 |
| Ulrich(H) | 0.98 | 0.96 | 0.97 | 0.98 | 0.97 | | |
| Li | 0.99 | 0.99 | 0.99 | 0.99 | 0.99 | | |
| Li(H) | 0.99 | 0.96 | 0.99 | 0.99 | 0.99 | | |
| Qiu(L) | 0.98 | 0.82 | 0.98 | 0.97 | 0.95 | | |
| Ulrich | 0.98 | 0.97 | 0.98 | 0.98 | 0.97 | | |
| Qiu(E) | 0.98 | 0.97 | 0.98 | 0.97 | 0.98 | | |
| Ayhan | 0.96 | 0.51 | 0.95 | 0.93 | 0.94 | | |
| Posner | 0.99 | 0.95 | 0.99 | 0.99 | 0.99 | | |
| Li(M) | 1.00 | 0.99 | 1.00 | 1.00 | 1.00 | | |
| Hoo | 1.00 | 1.00 | 1.00 | 1.00 | 1.00 | | |
| Yu | 0.98 | 0.98 | 0.99 | 0.98 | 0.99 | | |
| Yao | 1.00 | 1.00 | 1.00 | 1.00 | 1.00 | | |
| Li(HA) | 0.99 | 0.97 | 0.99 | 0.94 | 0.99 | | |
| Average | 0.98 | 0.92 | 0.94 | 0.96 | 0.95 | 0.94 | 0.94 |

Table S8: Gene Selection Algorithm Performance on 36 Benchmark scRNA-seq Datasets, Measured by Cell-type Local Inverse Simpson Index

| Data Name | Cell-type Local Inverse Simpson Index | | | | | | |
|---|---|---|---|---|---|---|---|
| | CellScope | Scanpy | Disp | VST | Mixhvg | FEAST | HRG |
| Yan | 1.22 | 1.35 | 1.22 | 1.24 | 1.23 | 1.40 | 1.39 |
| Goolam | 1.14 | 1.17 | 1.18 | 1.16 | 1.15 | 1.22 | 1.17 |
| Pollen | 1.17 | 1.17 | 1.18 | 1.16 | 1.22 | 1.14 | 1.14 |
| Wang | 1.08 | 1.18 | 1.13 | 1.17 | 1.12 | 1.09 | 1.21 |
| Darmanis | 1.23 | 1.67 | 1.58 | 1.38 | 1.64 | 1.98 | 1.42 |
| Usoskin | 1.06 | 1.39 | 1.34 | 1.40 | 1.26 | 1.83 | 1.97 |
| Xin | 1.15 | 1.28 | 1.23 | 1.15 | 1.29 | 1.25 | 2.04 |
| Keller(I) | 1.23 | 1.04 | 3.38 | 1.32 | 1.82 | 1.49 | 1.63 |
| Muraro | 1.06 | 1.08 | 1.08 | 1.07 | 1.06 | 1.34 | 1.22 |
| NHGRI | 1.05 | 1.10 | 1.10 | 1.10 | 1.08 | 1.13 | 1.09 |
| Klein | 1.03 | 1.17 | 1.18 | 1.08 | 1.15 | 1.27 | 1.09 |
| Zeisel | 1.13 | 1.30 | 1.17 | 1.15 | 1.15 | 1.26 | 1.16 |
| Lake | 1.22 | 2.43 | 1.79 | 1.81 | 1.76 | 1.26 | 1.59 |
| Keller(P) | 1.10 | 1.35 | 2.80 | 2.09 | 2.73 | 1.47 | 1.27 |
| Han(B) | 1.12 | 1.13 | 1.08 | 1.10 | 1.08 | | 1.09 |
| Siletti-1 | 1.00 | 1.00 | 1.01 | 1.01 | 1.01 | | 1.00 |
| Tirosh | 1.04 | 1.11 | 1.30 | 1.03 | 1.38 | | 1.36 |
| Siletti-2 | 1.00 | 1.01 | 1.00 | 1.00 | 1.00 | | 1.00 |
| Baron(H) | 1.03 | 1.06 | 1.03 | 1.03 | 1.04 | | 1.06 |
| Siletti-3 | 1.00 | 1.00 | 1.00 | 1.00 | 1.00 | | 1.00 |
| Siletti-4 | 1.00 | 1.02 | 1.00 | 1.00 | 1.00 | | 1.00 |
| Siletti-5 | 1.00 | 1.02 | 1.00 | 1.00 | 1.00 | | 1.00 |
| Siletti-6 | 1.00 | 1.04 | 1.00 | 1.00 | 1.00 | | 1.00 |
| Ulrich(H) | 1.06 | 1.07 | 1.06 | 1.05 | 1.07 | | |
| Li | 1.04 | 1.04 | 1.04 | 1.04 | 1.04 | | |
| Li(H) | 1.04 | 1.19 | 1.03 | 1.03 | 1.05 | | |
| Qiu(L) | 1.07 | 1.25 | 1.10 | 1.09 | 1.19 | | |
| Ulrich | 1.07 | 1.07 | 1.06 | 1.05 | 1.07 | | |
| Qiu(E) | 1.09 | 1.12 | 1.09 | 1.09 | 1.11 | | |
| Ayhan | 1.09 | 1.37 | 1.06 | 1.08 | 1.15 | | |
| Posner | 1.04 | 1.12 | 1.04 | 1.03 | 1.05 | | |
| Li(M) | 1.01 | 1.05 | 1.01 | 1.02 | 1.02 | | |
| Hoo | 1.02 | 1.02 | 1.02 | 1.02 | 1.02 | | |
| Yu | 1.03 | 1.02 | 1.03 | 1.03 | 1.04 | | |
| Yao | 1.00 | 1.00 | 1.00 | 1.00 | 1.00 | | |
| Li(HA) | 1.01 | 1.10 | 1.02 | 1.07 | 1.02 | | |
| Average | 1.07 | 1.18 | 1.23 | 1.14 | 1.19 | 1.37 | 1.26 |

Table S9: CellScope Graph-based Clustering Performance on 36 Benchmark scRNA-seq Datasets Using Different Gene Selection Methods, Measured by ARI

| Data Name | CellScope | Scanpy | Disp | VST | Mixhvg | FEAST | HRG |
|---|---|---|---|---|---|---|---|
| Yan | 0.97 | 0.90 | 0.84 | 0.86 | 0.85 | 0.83 | 0.90 |
| Goolam | 0.91 | 0.88 | 0.55 | 0.91 | 0.91 | 0.88 | 0.88 |
| Pollen | 0.90 | 0.75 | 0.78 | 0.81 | 0.88 | 0.91 | 0.90 |
| Wang | 0.86 | 0.78 | 0.83 | 0.77 | 0.82 | 0.87 | 0.72 |
| Darmanis | 0.76 | 0.49 | 0.69 | 0.76 | 0.39 | 0.56 | 0.62 |
| Usoskin | 0.97 | 0.64 | 0.76 | 0.58 | 0.91 | 0.48 | 0.19 |
| Xin | 0.92 | 0.86 | 0.93 | 0.78 | 0.89 | 0.80 | 0.01 |
| Keller(I) | 0.80 | 0.18 | 0.07 | 0.27 | 0.45 | 0.62 | 0.24 |
| Muraro | 0.92 | 0.90 | 0.92 | 0.91 | 0.93 | 0.81 | 0.74 |
| NHGRI | 0.95 | 0.87 | 0.89 | 0.89 | 0.90 | 0.88 | 0.90 |
| Klein | 0.97 | 0.54 | 0.70 | 0.88 | 0.79 | 0.77 | 0.88 |
| Zeisel | 0.80 | 0.75 | 0.72 | 0.73 | 0.75 | 0.68 | 0.77 |
| Lake | 0.82 | 0.36 | 0.69 | 0.61 | 0.59 | 0.85 | 0.57 |
| Keller(P) | 0.88 | 0.09 | 0.11 | 0.03 | 0.28 | 0.41 | 0.80 |
| Han(B) | 0.89 | 0.87 | 0.91 | 0.61 | 0.91 | | 0.89 |
| Siletti-1 | 0.93 | 0.99 | 0.99 | 0.95 | 0.98 | | 0.94 |
| Tirosh | 0.90 | 0.62 | 0.87 | 0.69 | 0.84 | | 0.61 |
| Siletti-2 | 0.99 | 0.87 | 0.88 | 0.88 | 0.88 | | 0.88 |
| Baron(H) | 0.92 | 0.91 | 0.92 | 0.91 | 0.91 | | 0.92 |
| Siletti-3 | 0.84 | 0.84 | 0.84 | 0.91 | 0.84 | | 0.88 |
| Siletti-4 | 0.99 | 0.85 | 0.88 | 0.66 | 0.85 | | 0.93 |
| Siletti-5 | 0.88 | 0.69 | 0.85 | 0.56 | 0.73 | | 0.80 |
| Siletti-6 | 0.90 | 0.87 | 0.88 | 0.84 | 0.88 | | 0.94 |
| Ulrich(H) | 0.80 | 0.80 | 0.75 | 0.75 | 0.71 | | |
| Li | 0.86 | 0.86 | 0.84 | 0.87 | 0.83 | | |
| Li(H) | 0.95 | 0.96 | 0.96 | 0.97 | 0.97 | | |
| Qiu(L) | 0.75 | 0.55 | 0.65 | 0.60 | 0.66 | | |
| Ulrich | 0.79 | 0.76 | 0.78 | 0.80 | 0.70 | | |
| Qiu(E) | 0.81 | 0.78 | 0.74 | 0.68 | 0.72 | | |
| Ayhan | 0.88 | 0.80 | 0.87 | 0.86 | 0.85 | | |
| Posner | 0.82 | 0.77 | 0.83 | 0.74 | 0.84 | | |
| Li(M) | 0.80 | 0.94 | 0.89 | 0.97 | 0.90 | | |
| Hoo | 0.96 | 0.93 | 0.92 | 0.97 | 0.93 | | |
| Yu | 0.86 | 0.76 | 0.52 | 0.87 | 0.53 | | |
| Yao | 0.83 | 0.74 | 0.66 | 0.83 | 0.66 | | |
| Li(HA) | 0.88 | 0.81 | 0.87 | 0.86 | 0.89 | | |
| Average | 0.88 | 0.75 | 0.77 | 0.77 | 0.79 | 0.74 | 0.74 |

#### 2.6 Cell-type Detection Performance Analysis

Table S10: Performance Comparison on 36 Benchmark scRNA-seq Datasets, Measured by Silhouette Coefficients and F1 Scores for Rare and Non-rare Cell Types

| Data Name | F1 Score of rare cell types | | | F1 Score of non-rare cell types | | | Silhouette coefficients | | |
|---|---|---|---|---|---|---|---|---|---|
| | CellScope | Scanpy | Seurat | CellScope | Scanpy | Seurat | CellScope | Scanpy | Seurat |
| Yan | 0.00 | 0.00 | 0.00 | 1.00 | 0.85 | 0.99 | 0.82 | 0.59 | 0.67 |
| Goolam | 0.33 | 0.18 | 0.00 | 0.83 | 0.83 | 0.87 | 0.70 | 0.26 | 0.35 |
| Pollen | 0.60 | 0.00 | 0.00 | 0.00 | 0.00 | 0.00 | 0.72 | 0.59 | 0.75 |
| Wang | 0.00 | 0.25 | 0.00 | 0.95 | 0.94 | 0.89 | 0.18 | 0.25 | 0.14 |
| Darmanis | 0.79 | 0.25 | 0.37 | 0.92 | 0.88 | 0.71 | 0.46 | 0.20 | 0.22 |
| Usoskin | 0.00 | 0.00 | 0.00 | 0.99 | 0.92 | 0.93 | 0.77 | 0.46 | 0.51 |
| Xin | 0.18 | 0.19 | 0.14 | 0.97 | 0.99 | 0.99 | 0.44 | 0.24 | 0.51 |
| Keller(I) | 0.96 | 0.00 | 0.94 | 0.97 | 0.93 | 0.96 | 0.40 | 0.18 | 0.28 |
| Muraro | 0.59 | 0.57 | 0.17 | 0.97 | 0.97 | 0.89 | 0.38 | 0.47 | 0.13 |
| NHGRI | 0.00 | 0.00 | 0.00 | 0.98 | 0.95 | 0.91 | 0.72 | 0.41 | 0.44 |
| Klein | 0.00 | 0.00 | 0.00 | 0.99 | 0.95 | 0.96 | 0.74 | 0.73 | 0.72 |
| Zeisel | 0.77 | 0.71 | 0.77 | 0.93 | 0.90 | 0.90 | 0.50 | 0.31 | 0.39 |
| Lake | 0.56 | 0.60 | 0.23 | 0.96 | 0.92 | 0.94 | 0.41 | 0.28 | 0.10 |
| Keller(P) | 0.58 | 0.60 | 0.31 | 0.97 | 0.96 | 0.42 | 0.56 | -0.11 | -0.20 |
| Han(B) | 0.25 | 0.01 | 0.24 | 0.98 | 0.71 | 0.98 | 0.06 | -0.50 | 0.31 |
| Siletti-1 | 0.42 | 0.57 | 0.12 | 0.96 | 1.00 | 0.96 | 0.72 | 0.19 | 0.47 |
| Tirosh | 0.27 | 0.28 | 0.42 | 0.95 | 0.94 | 0.95 | 0.44 | 0.32 | 0.39 |
| Siletti-2 | 0.33 | 0.17 | 0.33 | 1.00 | 0.99 | 0.89 | 0.69 | 0.42 | 0.57 |
| Baron(H) | 0.53 | 0.50 | 0.28 | 0.98 | 0.99 | 0.98 | 0.60 | 0.42 | 0.36 |
| Siletti-3 | 0.69 | 0.63 | 0.43 | 0.97 | 0.97 | 0.97 | 0.54 | 0.30 | 0.34 |
| Siletti-4 | 0.25 | 0.48 | 0.23 | 1.00 | 0.85 | 0.85 | 0.52 | 0.07 | 0.09 |
| Siletti-5 | 0.10 | 0.29 | 0.06 | 0.98 | 0.84 | 0.72 | 0.26 | -0.26 | -0.10 |
| Siletti-6 | 0.14 | 0.10 | 0.35 | 0.98 | 0.95 | 0.85 | 0.50 | 0.11 | 0.15 |
| Ulrich(H) | 0.76 | 0.39 | 0.78 | 0.98 | 0.98 | 0.98 | 0.59 | 0.25 | 0.48 |
| Li | 0.00 | 0.00 | 0.00 | 0.98 | 0.91 | 0.99 | 0.49 | 0.30 | 0.23 |
| Li(H) | 0.69 | 0.37 | 0.54 | 0.99 | 0.99 | 0.99 | 0.72 | 0.34 | 0.48 |
| Qiu(L) | 0.00 | 0.01 | 0.00 | 0.92 | 0.85 | 0.93 | 0.42 | 0.32 | 0.40 |
| Ulrich | 0.96 | 0.96 | 0.78 | 0.98 | 0.90 | 0.98 | 0.59 | 0.28 | 0.49 |
| Qiu(E) | 0.48 | 0.00 | 0.16 | 0.96 | 0.97 | 0.96 | 0.12 | 0.14 | 0.18 |
| Ayhan | 0.30 | 0.00 | 0.23 | 0.97 | 0.97 | 0.98 | 0.57 | 0.35 | 0.47 |
| Posner | 0.33 | 0.22 | 0.06 | 0.99 | 1.00 | 0.93 | 0.38 | 0.08 | 0.15 |
| Li(M) | 0.27 | 0.21 | 0.13 | 0.88 | 0.88 | 1.00 | 0.49 | 0.22 | 0.36 |
| Hoo | 0.88 | 0.95 | 0.95 | 0.99 | 0.96 | 0.99 | 0.70 | 0.46 | 0.49 |
| Yu | 0.22 | 0.03 | 0.19 | 0.96 | 0.96 | 0.95 | 0.14 | -0.06 | 0.07 |
| Yao | 0.17 | 0.27 | 0.10 | 0.97 | 0.82 | 0.86 | 0.53 | 0.15 | 0.29 |
| Li(HA) | 0.21 | 0.23 | 0.30 | 0.88 | 0.96 | 0.98 | 0.44 | 0.21 | 0.31 |
| Average | 0.38 | 0.28 | 0.27 | 0.94 | 0.90 | 0.89 | 0.51 | 0.25 | 0.33 |

#### 2.7 Parameter Sensitivity Analysis

Table S11: Clustering Performance on small-scaled Benchmark scRNA-seq Datasets, Measured by ARI under Different Parameters (PCA Dimensions, Number of Selected Genes, Number of Manifold Seeds)

| Data Name | PCA dimensions | | | | | | | Numbers of selected genes | | | | | |
|---|---|---|---|---|---|---|---|---|---|---|---|---|---|
| | 2 | 50 | 100 | 200 | 300 | 400 | 500 | 50 | 100 | 500 | 1000 | 5000 | 10000 |
| Yan | 0.92 | 0.84 | 0.97 | 0.97 | 0.97 | 0.97 | 0.97 | 0.80 | 0.83 | 0.97 | 0.95 | 0.90 | 0.84 |
| Goolam | 0.91 | 0.55 | 0.91 | 0.47 | 0.47 | 0.47 | 0.47 | 0.88 | 0.91 | 0.91 | 0.88 | 0.88 | 0.88 |
| Pollen | 0.83 | 0.90 | 0.90 | 0.91 | 0.90 | 0.90 | 0.90 | 0.87 | 0.92 | 0.90 | 0.90 | 0.86 | 0.84 |
| Wang | 0.86 | 0.90 | 0.86 | 0.89 | 0.91 | 0.76 | 0.78 | 0.79 | 0.86 | 0.86 | 0.86 | 0.84 | 0.84 |
| Darmanis | 0.79 | 0.79 | 0.76 | 0.64 | 0.43 | 0.55 | 0.64 | 0.66 | 0.71 | 0.76 | 0.80 | 0.79 | 0.76 |
| Usoskin | 0.34 | 0.97 | 0.97 | 0.97 | 0.94 | 0.94 | 0.96 | 0.85 | 0.93 | 0.97 | 0.93 | 0.86 | 0.79 |
| Xin | 0.90 | 0.92 | 0.92 | 0.95 | 0.94 | 0.88 | 0.91 | 0.19 | 0.26 | 0.92 | 0.94 | 0.90 | 0.92 |
| Keller(I) | 0.81 | 0.80 | 0.80 | 0.81 | 0.68 | 0.77 | 0.64 | 0.49 | 0.69 | 0.80 | 0.72 | 0.79 | 0.66 |
| Muraro | 0.91 | 0.91 | 0.92 | 0.89 | 0.89 | 0.89 | 0.89 | 0.87 | 0.94 | 0.92 | 0.95 | 0.90 | 0.92 |
| NHGRI | 0.94 | 0.95 | 0.95 | 0.94 | 0.94 | 0.93 | 0.94 | 0.90 | 0.94 | 0.95 | 0.94 | 0.87 | 0.89 |
| Klein | 0.97 | 0.83 | 0.97 | 0.96 | 0.96 | 0.90 | 0.95 | 0.83 | 0.93 | 0.97 | 0.96 | 0.94 | 0.96 |
| Zeisel | 0.77 | 0.78 | 0.80 | 0.73 | 0.77 | 0.76 | 0.80 | 0.66 | 0.76 | 0.80 | 0.82 | 0.71 | 0.70 |
| Lake | 0.77 | 0.86 | 0.82 | 0.76 | 0.80 | 0.84 | 0.84 | 0.71 | 0.84 | 0.82 | 0.80 | 0.58 | 0.45 |
| Keller(P) | 0.84 | 0.88 | 0.88 | 0.80 | 0.84 | 0.84 | 0.84 | 0.73 | 0.84 | 0.88 | 0.88 | 0.65 | 0.55 |
| Han(B) | 0.90 | 0.91 | 0.89 | 0.78 | 0.80 | 0.80 | 0.80 | 0.56 | 0.64 | 0.89 | 0.88 | 0.88 | 0.79 |
| Siletti-1 | 0.93 | 0.93 | 0.93 | 0.93 | 0.93 | 0.93 | 0.93 | 1.00 | 1.00 | 0.93 | 0.93 | 0.93 | 0.93 |
| Tirosh | 0.63 | 0.61 | 0.90 | 0.61 | 0.54 | 0.53 | 0.61 | 0.82 | 0.88 | 0.90 | 0.82 | 0.55 | 0.57 |
| Siletti-2 | 0.88 | 1.00 | 0.99 | 1.00 | 0.99 | 0.99 | 0.99 | 0.99 | 0.99 | 0.99 | 1.00 | 1.00 | 1.00 |
| Baron(H) | 0.92 | 0.94 | 0.92 | 0.93 | 0.87 | 0.74 | 0.86 | 0.91 | 0.92 | 0.92 | 0.93 | 0.94 | 0.89 |
| Siletti-3 | 0.91 | 0.84 | 0.84 | 0.84 | 0.84 | 0.99 | 0.84 | 0.72 | 0.87 | 0.84 | 0.85 | 0.84 | 0.84 |
| Siletti-4 | 0.92 | 0.72 | 0.99 | 0.99 | 0.88 | 0.88 | 0.88 | 0.95 | 0.96 | 0.99 | 0.99 | 0.96 | 0.89 |
| Siletti-5 | 0.68 | 0.74 | 0.88 | 0.47 | 0.62 | 0.65 | 0.74 | 0.89 | 0.38 | 0.88 | 0.76 | 0.84 | 0.80 |
| Siletti-6 | 0.89 | 0.88 | 0.90 | 0.88 | 0.88 | 0.91 | 0.91 | 0.89 | 0.92 | 0.90 | 0.88 | 0.89 | 0.91 |
| Average | 0.84 | 0.84 | 0.90 | 0.83 | 0.82 | 0.82 | 0.83 | 0.78 | 0.82 | 0.90 | 0.89 | 0.84 | 0.81 |

| Data Name | Numbers of manifold seeds during gene selection | | | | | | | | | | | | | | |
|---|---|---|---|---|---|---|---|---|---|---|---|---|---|---|---|
| | 3 | 4 | 5 | 6 | 7 | 8 | 9 | 10 | 11 | 12 | 13 | 14 | 15 | 16 | Adaptive |
| Yan | 0.97 | 0.86 | 0.90 | 0.80 | 0.80 | 0.79 | 0.81 | 0.84 | 0.92 | 0.88 | 0.87 | 0.88 | 0.92 | 0.89 | 0.97 |
| Goolam | 0.91 | 0.99 | 0.88 | 0.98 | 0.98 | 0.99 | 0.99 | 0.88 | 0.91 | 0.99 | 0.99 | 0.99 | 0.99 | 0.99 | 0.91 |
| Pollen | 0.91 | 0.90 | 0.88 | 0.89 | 0.90 | 0.88 | 0.85 | 0.81 | 0.83 | 0.74 | 0.74 | 0.76 | 0.74 | 0.75 | 0.90 |
| Wang | 0.86 | 0.89 | 0.91 | 0.88 | 0.83 | 0.89 | 0.82 | 0.81 | 0.82 | 0.89 | 0.86 | 0.86 | 0.83 | 0.84 | 0.86 |
| Darmanis | 0.76 | 0.76 | 0.77 | 0.75 | 0.74 | 0.75 | 0.80 | 0.81 | 0.77 | 0.74 | 0.73 | 0.75 | 0.68 | 0.65 | 0.76 |
| Usoskin | 0.97 | 0.98 | 0.95 | 0.90 | 0.87 | 0.65 | 0.84 | 0.73 | 0.60 | 0.87 | 0.63 | 0.47 | 0.45 | 0.38 | 0.97 |
| Xin | 0.92 | 0.91 | 0.93 | 0.97 | 0.94 | 0.93 | 0.95 | 0.91 | 0.91 | 0.87 | 0.91 | 0.73 | 0.73 | 0.70 | 0.92 |
| Keller(I) | 0.74 | 0.81 | 0.81 | 0.79 | 0.71 | 0.81 | 0.76 | 0.73 | 0.65 | 0.78 | 0.80 | 0.77 | 0.81 | 0.67 | 0.80 |
| Muraro | 0.89 | 0.89 | 0.90 | 0.91 | 0.91 | 0.85 | 0.91 | 0.91 | 0.90 | 0.91 | 0.85 | 0.85 | 0.91 | 0.87 | 0.92 |
| NHGRI | 0.92 | 0.92 | 0.92 | 0.92 | 0.94 | 0.95 | 0.94 | 0.94 | 0.94 | 0.94 | 0.94 | 0.90 | 0.93 | 0.94 | 0.95 |
| Klein | 0.65 | 0.94 | 0.95 | 0.97 | 0.94 | 0.88 | 0.97 | 0.98 | 0.96 | 0.97 | 0.85 | 0.97 | 0.89 | 0.86 | 0.97 |
| Zeisel | 0.75 | 0.77 | 0.74 | 0.76 | 0.77 | 0.77 | 0.76 | 0.79 | 0.76 | 0.75 | 0.78 | 0.78 | 0.79 | 0.80 | 0.80 |
| Lake | 0.82 | 0.83 | 0.78 | 0.80 | 0.82 | 0.85 | 0.78 | 0.83 | 0.83 | 0.85 | 0.84 | 0.84 | 0.81 | 0.85 | 0.82 |
| Keller(P) | 0.77 | 0.82 | 0.79 | 0.80 | 0.79 | 0.74 | 0.80 | 0.88 | 0.88 | 0.85 | 0.88 | 0.87 | 0.88 | 0.87 | 0.88 |
| Han(B) | 0.90 | 0.87 | 0.91 | 0.89 | 0.90 | 0.90 | 0.90 | 0.90 | 0.90 | 0.91 | 0.91 | 0.91 | 0.91 | 0.91 | 0.89 |
| Siletti-1 | 0.93 | 0.93 | 0.93 | 0.93 | 0.93 | 0.93 | 0.93 | 0.93 | 0.93 | 0.93 | 0.93 | 0.93 | 0.93 | 0.93 | 0.93 |
| Tirosh | 0.52 | 0.58 | 0.62 | 0.62 | 0.63 | 0.84 | 0.88 | 0.88 | 0.84 | 0.67 | 0.88 | 0.88 | 0.88 | 0.88 | 0.90 |
| Siletti-2 | 0.99 | 0.99 | 1.00 | 0.98 | 1.00 | 1.00 | 0.88 | 1.00 | 0.88 | 1.00 | 0.88 | 0.88 | 0.88 | 0.88 | 0.99 |
| Baron(H) | 0.69 | 0.81 | 0.85 | 0.89 | 0.87 | 0.90 | 0.86 | 0.86 | 0.86 | 0.93 | 0.92 | 0.91 | 0.89 | 0.93 | 0.92 |
| Siletti-3 | 0.84 | 0.84 | 0.84 | 0.84 | 0.84 | 0.84 | 0.87 | 0.84 | 0.84 | 0.96 | 0.98 | 0.84 | 0.97 | 0.84 | 0.84 |
| Siletti-4 | 0.98 | 0.92 | 0.96 | 0.99 | 0.93 | 0.97 | 0.91 | 0.93 | 1.00 | 0.83 | 0.98 | 0.95 | 0.96 | 0.87 | 0.99 |
| Siletti-5 | 0.74 | 0.74 | 0.65 | 0.46 | 0.44 | 0.74 | 0.53 | 0.69 | 0.44 | 0.73 | 0.74 | 0.73 | 0.38 | 0.37 | 0.88 |
| Siletti-6 | 0.89 | 0.87 | 0.82 | 0.90 | 0.88 | 0.90 | 0.91 | 0.97 | 0.91 | 0.99 | 0.88 | 0.70 | 0.75 | 0.64 | 0.90 |
| Average | 0.84 | 0.86 | 0.86 | 0.85 | 0.84 | 0.86 | 0.85 | 0.86 | 0.84 | 0.87 | 0.86 | 0.83 | 0.82 | 0.80 | 0.90 |

#### 2.8 Robustness Analysis

Table S12: Clustering Performance on 36 Benchmark scRNA-seq Datasets, Measured by ARI for Three Robustness Tests: Manifold Fitting, Distance Metrics (Euclidean vs. Jaccard), and Clustering Methods (Graph-based vs. Distance-based)

| Data Name | ARI | | ARI | | ARI | |
|---|---|---|---|---|---|---|
| | With MF | WO MF | Euclidean | Jaccard | Graph | Distance |
| Yan | 0.97 | 0.80 | 0.97 | 0.97 | 0.97 | 0.83 |
| Goolam | 0.91 | 0.88 | 0.91 | 0.88 | 0.91 | 0.95 |
| Pollen | 0.90 | 0.78 | 0.90 | 0.90 | 0.90 | 0.89 |
| Wang | 0.86 | 0.85 | 0.86 | 0.79 | 0.86 | 0.85 |
| Darmanis | 0.76 | 0.72 | 0.76 | 0.73 | 0.76 | 0.75 |
| Usoskin | 0.97 | 0.74 | 0.97 | 0.87 | 0.97 | 0.66 |
| Xin | 0.92 | 0.85 | 0.92 | 0.43 | 0.92 | 0.74 |
| Keller(I) | 0.80 | 0.80 | 0.80 | 0.41 | 0.80 | 0.50 |
| Muraro | 0.92 | 0.88 | 0.92 | 0.93 | 0.92 | 0.74 |
| NHGRI | 0.95 | 0.87 | 0.95 | 0.88 | 0.95 | 0.86 |
| Klein | 0.97 | 0.94 | 0.97 | 0.65 | 0.97 | 0.80 |
| Zeisel | 0.80 | 0.76 | 0.80 | 0.72 | 0.80 | 0.52 |
| Lake | 0.82 | 0.44 | 0.82 | 0.38 | 0.82 | 0.71 |
| Keller(P) | 0.88 | 0.77 | 0.88 | 0.83 | 0.88 | 0.55 |
| Han(B) | 0.89 | 0.76 | 0.89 | 0.78 | 0.89 | 0.76 |
| Siletti-1 | 0.93 | 0.93 | 0.93 | 0.97 | 0.93 | 1.00 |
| Tirosh | 0.90 | 0.64 | 0.90 | 0.86 | 0.90 | 0.86 |
| Siletti-2 | 0.99 | 0.88 | 0.99 | 0.99 | 0.99 | 0.97 |
| Baron(H) | 0.92 | 0.88 | 0.92 | 0.86 | 0.92 | 0.86 |
| Siletti-3 | 0.84 | 0.85 | 0.84 | 0.84 | 0.84 | 0.03 |
| Siletti-4 | 0.99 | 0.99 | 0.92 | 0.99 | 0.99 | |
| Siletti-5 | 0.88 | 0.68 | 0.55 | 0.88 | 0.88 | |
| Siletti-6 | 0.90 | 0.97 | 0.87 | 0.90 | 0.90 | |
| Ulrich(H) | 0.80 | 0.72 | 0.62 | 0.80 | 0.80 | |
| Li | 0.86 | 0.85 | 0.52 | 0.87 | 0.86 | |
| Li(H) | 0.95 | 0.87 | 0.95 | 0.95 | 0.95 | |
| Qiu(L) | 0.75 | 0.57 | 0.60 | 0.69 | 0.75 | |
| Ulrich | 0.79 | 0.76 | 0.62 | 0.79 | 0.79 | |
| Qiu(E) | 0.81 | 0.71 | 0.75 | 0.81 | 0.81 | |
| Ayhan | 0.88 | 0.89 | 0.84 | 0.89 | 0.88 | |
| Posner | 0.82 | 0.81 | 0.82 | 0.83 | 0.82 | |
| Li(M) | 0.80 | 0.88 | 0.81 | 0.81 | 0.80 | |
| Hoo | 0.96 | 0.97 | 0.92 | 0.97 | 0.96 | |
| Yu | 0.86 | 0.81 | 0.84 | 0.87 | 0.86 | |
| Yao | 0.83 | 0.97 | 0.94 | 0.81 | 0.83 | |
| Li(HA) | 0.88 | 0.87 | 0.91 | 0.88 | 0.88 | |
| Average | 0.88 | 0.82 | 0.84 | 0.82 | 0.88 | |

#### 2.9 Interaction analysis of gene selection

Table S13: High-Quality Gene Selection Analysis on 36 Benchmark scRNA-seq Datasets: Comparison of CellScope, Scanpy, and Seurat

| Data Name | CellScope | Scanpy | Seurat | CellScope/Scanpy | Scanpy/CellScope | CellScope/Seurat | Seurat/CellScope |
|---|---|---|---|---|---|---|---|
| Yan | 485 (97.0%) | 3149 (59.6%) | 1091 (54.5%) | 163 (93.7%) | 2827 (57.0%) | 281 (95.6%) | 887 (49.4%) |
| Goolam | 381 (76.2%) | 2208 (32.1%) | 1043 (52.1%) | 102 (66.7%) | 1929 (29.5%) | 204 (65.4%) | 866 (47.8%) |
| Pollen | 308 (61.6%) | 543 (13.0%) | 361 (18.1%) | 170 (60.5%) | 405 (10.3%) | 219 (61.5%) | 272 (14.7%) |
| Wang | 187 (37.4%) | 218 (3.0%) | 132 (6.6%) | 60 (31.7%) | 91 (1.3%) | 108 (28.8%) | 53 (2.8%) |
| Darmanis | 90 (18.0%) | 173 (2.6%) | 123 (6.2%) | 17 (9.0%) | 100 (1.6%) | 39 (10.5%) | 72 (3.8%) |
| Usoskin | 20 (4.0%) | 14 (0.3%) | 12 (0.6%) | 7 (2.1%) | 1 (0.0%) | 9 (2.2%) | 1 (0.1%) |
| Xin | 37 (7.4%) | 58 (0.7%) | 27 (1.4%) | 8 (3.9%) | 29 (0.3%) | 23 (5.6%) | 13 (0.7%) |
| Keller(I) | 179 (35.8%) | 86 (2.4%) | 38 (1.9%) | 135 (32.7%) | 42 (1.2%) | 157 (34.1%) | 16 (0.8%) |
| Muraro | 131 (26.2%) | 714 (32.5%) | 590 (29.5%) | 40 (12.4%) | 623 (30.8%) | 75 (18.6%) | 534 (28.1%) |
| NHGRI | 440 (88.0%) | 1106 (39.6%) | 468 (23.4%) | 200 (90.1%) | 866 (34.4%) | 289 (91.2%) | 317 (17.4%) |
| Klein | 77 (15.4%) | 53 (0.9%) | 130 (6.5%) | 59 (19.3%) | 35 (0.6%) | 61 (15.9%) | 114 (6.1%) |
| Zeisel | 141 (28.2%) | 172 (4.3%) | 178 (8.9%) | 80 (22.8%) | 111 (2.9%) | 76 (23.1%) | 113 (6.2%) |
| Lake | 135 (27.0%) | 133 (1.8%) | 55 (2.8%) | 38 (16.4%) | 36 (0.5%) | 98 (23.0%) | 18 (0.9%) |
| Keller(P) | 235 (47.0%) | 55 (1.1%) | 0 (0.0%) | 206 (47.8%) | 26 (0.5%) | 235 (47.0%) | 0 (0.0%) |
| Han(B) | 43 (8.6%) | 67 (2.4%) | 96 (4.8%) | 20 (5.8%) | 44 (1.7%) | 9 (3.0%) | 62 (3.4%) |
| Siletti-1 | 324 (64.8%) | 896 (16.7%) | 229 (11.5%) | 209 (59.9%) | 781 (15.0%) | 296 (64.3%) | 201 (10.3%) |
| Tirosh | 157 (31.4%) | 226 (4.5%) | 166 (8.3%) | 47 (26.6%) | 116 (2.5%) | 74 (23.8%) | 83 (4.6%) |
| Siletti-2 | 260 (52.0%) | 630 (8.5%) | 451 (22.6%) | 159 (47.3%) | 529 (7.3%) | 140 (46.1%) | 331 (18.3%) |
| Baron(H) | 195 (39.0%) | 212 (11.4%) | 224 (11.2%) | 37 (34.9%) | 54 (3.7%) | 25 (32.5%) | 54 (3.4%) |
| Siletti-3 | 311 (62.2%) | 658 (14.7%) | 512 (25.6%) | 105 (58.0%) | 452 (10.9%) | 143 (73.7%) | 344 (20.3%) |
| Siletti-4 | 168 (33.6%) | 443 (6.9%) | 276 (13.8%) | 103 (33.4%) | 378 (6.1%) | 120 (36.0%) | 228 (12.4%) |
| Siletti-5 | 298 (59.6%) | 576 (9.6%) | 504 (25.2%) | 158 (76.0%) | 436 (7.6%) | 157 (72.4%) | 363 (21.1%) |
| Siletti-6 | 309 (61.8%) | 647 (9.8%) | 429 (21.4%) | 149 (70.3%) | 487 (7.7%) | 209 (68.8%) | 329 (18.2%) |
| Ulrich(H) | 213 (42.6%) | 360 (11.1%) | 332 (16.6%) | 11 (10.2%) | 158 (5.5%) | 34 (28.1%) | 153 (9.4%) |
| Li | 26 (5.2%) | 29 (1.1%) | 26 (1.3%) | 0 (0.0%) | 3 (0.1%) | 3 (1.4%) | 3 (0.2%) |
| Li(H) | 54 (10.8%) | 54 (1.2%) | 56 (2.8%) | 3 (1.5%) | 3 (0.1%) | 1 (0.5%) | 3 (0.2%) |
| Qiu(L) | 20 (4.0%) | 19 (0.5%) | 19 (0.9%) | 1 (0.6%) | 0 (0.0%) | 1 (0.9%) | 0 (0.0%) |
| Ulrich | 219 (43.8%) | 376 (12.0%) | 351 (17.5%) | 11 (9.6%) | 168 (6.1%) | 36 (26.5%) | 168 (10.3%) |
| Qiu(E) | 71 (14.2%) | 91 (2.9%) | 76 (3.8%) | 17 (12.0%) | 37 (1.3%) | 35 (21.5%) | 40 (2.4%) |
| Ayhan | 15 (3.0%) | 8 (0.2%) | 19 (0.9%) | 8 (3.4%) | 1 (0.0%) | 0 (0.0%) | 4 (0.2%) |
| Posner | 182 (36.4%) | 195 (4.9%) | 204 (10.2%) | 48 (20.3%) | 61 (1.6%) | 59 (35.3%) | 81 (4.9%) |
| Li(M) | 82 (16.4%) | 101 (4.2%) | 96 (4.8%) | 3 (1.2%) | 22 (1.0%) | 8 (3.2%) | 22 (1.3%) |
| Hoo | 157 (31.4%) | 364 (7.5%) | 162 (8.1%) | 16 (19.0%) | 223 (5.0%) | 87 (29.1%) | 92 (5.1%) |
| Yu | 187 (37.4%) | 371 (15.4%) | 411 (20.5%) | 19 (21.6%) | 203 (10.2%) | 32 (29.1%) | 256 (15.9%) |
| Yao | 344 (68.8%) | 542 (12.6%) | 498 (24.9%) | 125 (71.4%) | 323 (8.1%) | 140 (80.5%) | 294 (17.6%) |
| Li(HA) | 191 (38.2%) | 219 (4.3%) | 228 (11.4%) | 17 (13.7%) | 45 (0.9%) | 90 (39.3%) | 127 (7.3%) |

Table S14: Low-Quality Gene Selection Analysis on 36 Benchmark scRNA-seq Datasets: Comparison of CellScope, Scanpy, and Seurat

| Data Name | CellScope | Scanpy | Seurat | CellScope/Scanpy | Scanpy/CellScope | CellScope/Seurat | Seurat/CellScope |
|---|---|---|---|---|---|---|---|
| Yan | 15 (3.0 %) | 2137 (40.4 %) | 909 (45.5 %) | 11 (6.3 %) | 2133 (43.0 %) | 13 (4.4 %) | 907 (50.6 %) |
| Goolam | 119 (23.8 %) | 4679 (67.9 %) | 957 (47.9 %) | 51 (33.3 %) | 4611 (70.5 %) | 108 (34.6 %) | 946 (52.2 %) |
| Pollen | 192 (38.4 %) | 3620 (87.0 %) | 1639 (82.0 %) | 111 (39.5 %) | 3539 (89.7 %) | 137 (38.5 %) | 1584 (85.3 %) |
| Wang | 313 (62.6 %) | 7125 (97.0 %) | 1868 (93.4 %) | 129 (68.3 %) | 6941 (98.7 %) | 267 (71.2 %) | 1822 (97.2 %) |
| Darmanis | 410 (82.0 %) | 6522 (97.4 %) | 1877 (93.8 %) | 172 (91.0 %) | 6284 (98.4 %) | 334 (89.5 %) | 1801 (96.2 %) |
| Usoskin | 480 (96.0 %) | 4170 (99.7 %) | 1988 (99.4 %) | 334 (97.9 %) | 4024 (100.0 %) | 395 (97.8 %) | 1903 (99.9 %) |
| Xin | 463 (92.6 %) | 8682 (99.3 %) | 1973 (98.7 %) | 195 (96.1 %) | 8414 (99.7 %) | 389 (94.4 %) | 1899 (99.3 %) |
| Keller(I) | 321 (64.2 %) | 3559 (97.6 %) | 1962 (98.1 %) | 278 (67.3 %) | 3516 (98.8 %) | 304 (65.9 %) | 1945 (99.2 %) |
| Muraro | 369 (73.8 %) | 1486 (67.5 %) | 1410 (70.5 %) | 283 (87.6 %) | 1400 (69.2 %) | 328 (81.4 %) | 1369 (71.9 %) |
| NHGRI | 60 (12.0 %) | 1686 (60.4 %) | 1532 (76.6 %) | 22 (9.9 %) | 1648 (65.6 %) | 28 (8.8 %) | 1500 (82.6 %) |
| Klein | 423 (84.6 %) | 5654 (99.1 %) | 1870 (93.5 %) | 247 (80.7 %) | 5478 (99.4 %) | 323 (84.1 %) | 1770 (93.9 %) |
| Zeisel | 359 (71.8 %) | 3868 (95.7 %) | 1822 (91.1 %) | 271 (77.2 %) | 3780 (97.1 %) | 253 (76.9 %) | 1716 (93.8 %) |
| Lake | 365 (73.0 %) | 7280 (98.2 %) | 1945 (97.2 %) | 194 (83.6 %) | 7109 (99.5 %) | 328 (77.0 %) | 1908 (99.1 %) |
| Keller(P) | 265 (53.0 %) | 5015 (98.9 %) | 2000 (100.0 %) | 225 (52.2 %) | 4975 (99.5 %) | 265 (53.0 %) | 2000 (100.0 %) |
| Han(B) | 457 (91.4 %) | 2739 (97.6 %) | 1904 (95.2 %) | 323 (94.2 %) | 2605 (98.3 %) | 294 (97.0 %) | 1741 (96.6 %) |
| Siletti-1 | 176 (35.2 %) | 4479 (83.3 %) | 1771 (88.5 %) | 140 (40.1 %) | 4443 (85.0 %) | 164 (35.7 %) | 1759 (89.7 %) |
| Tirosh | 343 (68.6 %) | 4792 (95.5 %) | 1834 (91.7 %) | 130 (73.4 %) | 4579 (97.5 %) | 237 (76.2 %) | 1728 (95.4 %) |
| Siletti-2 | 240 (48.0 %) | 6740 (91.5 %) | 1549 (77.5 %) | 177 (52.7 %) | 6677 (92.7 %) | 164 (53.9 %) | 1473 (81.7 %) |
| Baron(H) | 305 (61.0 %) | 1652 (88.6 %) | 1776 (88.8 %) | 69 (65.1 %) | 1416 (96.3 %) | 52 (67.5 %) | 1523 (96.6 %) |
| Siletti-3 | 189 (37.8 %) | 3815 (85.3 %) | 1488 (74.4 %) | 76 (42.0 %) | 3702 (89.1 %) | 51 (26.3 %) | 1350 (79.7 %) |
| Siletti-4 | 332 (66.4 %) | 5937 (93.1 %) | 1724 (86.2 %) | 205 (66.6 %) | 5810 (93.9 %) | 213 (64.0 %) | 1605 (87.6 %) |
| Siletti-5 | 202 (40.4 %) | 5420 (90.4 %) | 1496 (74.8 %) | 50 (24.0 %) | 5268 (92.4 %) | 60 (27.6 %) | 1354 (78.9 %) |
| Siletti-6 | 191 (38.2 %) | 5959 (90.2 %) | 1571 (78.5 %) | 63 (29.7 %) | 5831 (92.3 %) | 95 (31.2 %) | 1475 (81.8 %) |
| Ulrich(H) | 287 (57.4 %) | 2883 (88.9 %) | 1668 (83.4 %) | 97 (89.8 %) | 2693 (94.5 %) | 87 (71.9 %) | 1468 (90.6 %) |
| Li | 474 (94.8 %) | 2607 (98.9 %) | 1974 (98.7 %) | 178 (100.0 %) | 2311 (99.9 %) | 206 (98.6 %) | 1706 (99.8 %) |
| Li(H) | 446 (89.2 %) | 4443 (98.8 %) | 1944 (97.2 %) | 200 (98.5 %) | 4197 (99.9 %) | 182 (99.5 %) | 1680 (99.8 %) |
| Qiu(L) | 480 (96.0 %) | 3994 (99.5 %) | 1981 (99.1 %) | 174 (99.4 %) | 3688 (100.0 %) | 105 (99.1 %) | 1606 (100.0 %) |
| Ulrich | 281 (56.2 %) | 2766 (88.0 %) | 1649 (82.5 %) | 104 (90.4 %) | 2589 (93.9 %) | 100 (73.5 %) | 1468 (89.7 %) |
| Qiu(E) | 429 (85.8 %) | 3019 (97.1 %) | 1924 (96.2 %) | 125 (88.0 %) | 2715 (98.7 %) | 128 (78.5 %) | 1623 (97.6 %) |
| Ayhan | 485 (97.0 %) | 3422 (99.8 %) | 1981 (99.1 %) | 229 (96.6 %) | 3166 (100.0 %) | 251 (100.0 %) | 1747 (99.8 %) |
| Posner | 318 (63.6 %) | 3770 (95.1 %) | 1796 (89.8 %) | 188 (79.7 %) | 3640 (98.4 %) | 108 (64.7 %) | 1586 (95.1 %) |
| Li(M) | 418 (83.6 %) | 2316 (95.8 %) | 1904 (95.2 %) | 249 (98.8 %) | 2147 (99.0 %) | 245 (96.8 %) | 1731 (98.7 %) |
| Hoo | 343 (68.6 %) | 4488 (92.5 %) | 1838 (91.9 %) | 68 (81.0 %) | 4213 (95.0 %) | 212 (70.9 %) | 1707 (94.9 %) |
| Yu | 313 (62.6 %) | 2035 (84.6 %) | 1589 (79.5 %) | 69 (78.4 %) | 1791 (89.8 %) | 78 (70.9 %) | 1354 (84.1 %) |
| Yao | 156 (31.2 %) | 3748 (87.4 %) | 1502 (75.1 %) | 50 (28.6 %) | 3642 (91.9 %) | 34 (19.5 %) | 1380 (82.4 %) |
| Li(HA) | 309 (61.8 %) | 4917 (95.7 %) | 1772 (88.6 %) | 107 (86.3 %) | 4715 (99.1 %) | 139 (60.7 %) | 1602 (92.7 %) |

Table S15: Ratios of High-Quality to All Genes Selected by CellScope, Scanpy, and Seurat Across 36 Benchmark Datasets, with Comparisons Excluding Intersections (CellScope/Scanpy, Scanpy/CellScope, CellScope/Seurat, Seurat/CellScope)

| Data Name | High Quality Genes / All Genes (%) | | | | | | |
|---|---|---|---|---|---|---|---|
| | CellScope | Scanpy | Seurat | CellScope/Scanpy | Scanpy/CellScope | CellScope/Seurat | Seurat/CellScope |
| Yan | 97.00% | 59.60% | 54.50% | 93.70% | 57.00% | 95.60% | 49.40% |
| Goolam | 76.20% | 32.10% | 52.10% | 66.70% | 29.50% | 65.40% | 47.80% |
| Pollen | 61.60% | 13.00% | 18.10% | 60.50% | 10.30% | 61.50% | 14.70% |
| Wang | 37.40% | 3.00% | 6.60% | 31.70% | 1.30% | 28.80% | 2.80% |
| Darmanis | 18.00% | 2.60% | 6.20% | 9.00% | 1.60% | 10.50% | 3.80% |
| Usoskin | 4.00% | 0.30% | 0.60% | 2.10% | 0.00% | 2.20% | 0.10% |
| Xin | 7.40% | 0.70% | 1.40% | 3.90% | 0.30% | 5.60% | 0.70% |
| Keller(I) | 35.80% | 2.40% | 1.90% | 32.70% | 1.20% | 34.10% | 0.80% |
| Muraro | 26.20% | 32.50% | 29.50% | 12.40% | 30.80% | 18.60% | 28.10% |
| NHGRI | 88.00% | 39.60% | 23.40% | 90.10% | 34.40% | 91.20% | 17.40% |
| Klein | 15.40% | 0.90% | 6.50% | 19.30% | 0.60% | 15.90% | 6.10% |
| Zeisel | 28.20% | 4.30% | 8.90% | 22.80% | 2.90% | 23.10% | 6.20% |
| Lake | 27.00% | 1.80% | 2.80% | 16.40% | 0.50% | 23.00% | 0.90% |
| Keller(P) | 47.00% | 1.10% | 0.00% | 47.80% | 0.50% | 47.00% | 0.00% |
| Han(B) | 8.60% | 2.40% | 4.80% | 5.80% | 1.70% | 3.00% | 3.40% |
| Siletti-1 | 64.80% | 16.70% | 11.50% | 59.90% | 15.00% | 64.30% | 10.30% |
| Tirosh | 31.40% | 4.50% | 8.30% | 26.60% | 2.50% | 23.80% | 4.60% |
| Siletti-2 | 52.00% | 8.50% | 22.60% | 47.30% | 7.30% | 46.10% | 18.30% |
| Baron(H) | 39.00% | 11.40% | 11.20% | 34.90% | 3.70% | 32.50% | 3.40% |
| Siletti-3 | 62.20% | 14.70% | 25.60% | 58.00% | 10.90% | 73.70% | 20.30% |
| Siletti-4 | 33.60% | 6.90% | 13.80% | 33.40% | 6.10% | 36.00% | 12.40% |
| Siletti-5 | 59.60% | 9.60% | 25.20% | 76.00% | 7.60% | 72.40% | 21.10% |
| Siletti-6 | 61.80% | 9.80% | 21.40% | 70.30% | 7.70% | 68.80% | 18.20% |
| Ulrich(H) | 42.60% | 11.10% | 16.60% | 10.20% | 5.50% | 28.10% | 9.40% |
| Li | 5.20% | 1.10% | 1.30% | 0.00% | 0.10% | 1.40% | 0.20% |
| Li(H) | 10.80% | 1.20% | 2.80% | 1.50% | 0.10% | 0.50% | 0.20% |
| Qiu(L) | 4.00% | 0.50% | 0.90% | 0.60% | 0.00% | 0.90% | 0.00% |
| Ulrich | 43.80% | 12.00% | 17.50% | 9.60% | 6.10% | 26.50% | 10.30% |
| Qiu(E) | 14.20% | 2.90% | 3.80% | 12.00% | 1.30% | 21.50% | 2.40% |
| Ayhan | 3.00% | 0.20% | 0.90% | 3.40% | 0.00% | 0.00% | 0.20% |
| Posner | 36.40% | 4.90% | 10.20% | 20.30% | 1.60% | 35.30% | 4.90% |
| Li(M) | 16.40% | 4.20% | 4.80% | 1.20% | 1.00% | 3.20% | 1.30% |
| Hoo | 31.40% | 7.50% | 8.10% | 19.00% | 5.00% | 29.10% | 5.10% |
| Yu | 37.40% | 15.40% | 20.50% | 21.60% | 10.20% | 29.10% | 15.90% |
| Yao | 68.80% | 12.60% | 24.90% | 71.40% | 8.10% | 80.50% | 17.60% |
| Li(HA) | 38.20% | 4.30% | 11.40% | 13.70% | 0.90% | 39.30% | 7.30% |
| Average | 37.07% | 9.90% | 13.35% | 30.72% | 7.59% | 34.40% | 10.16% |

### 3 Supplementary Figure

#### 3.1 Clustering Performance Comparison

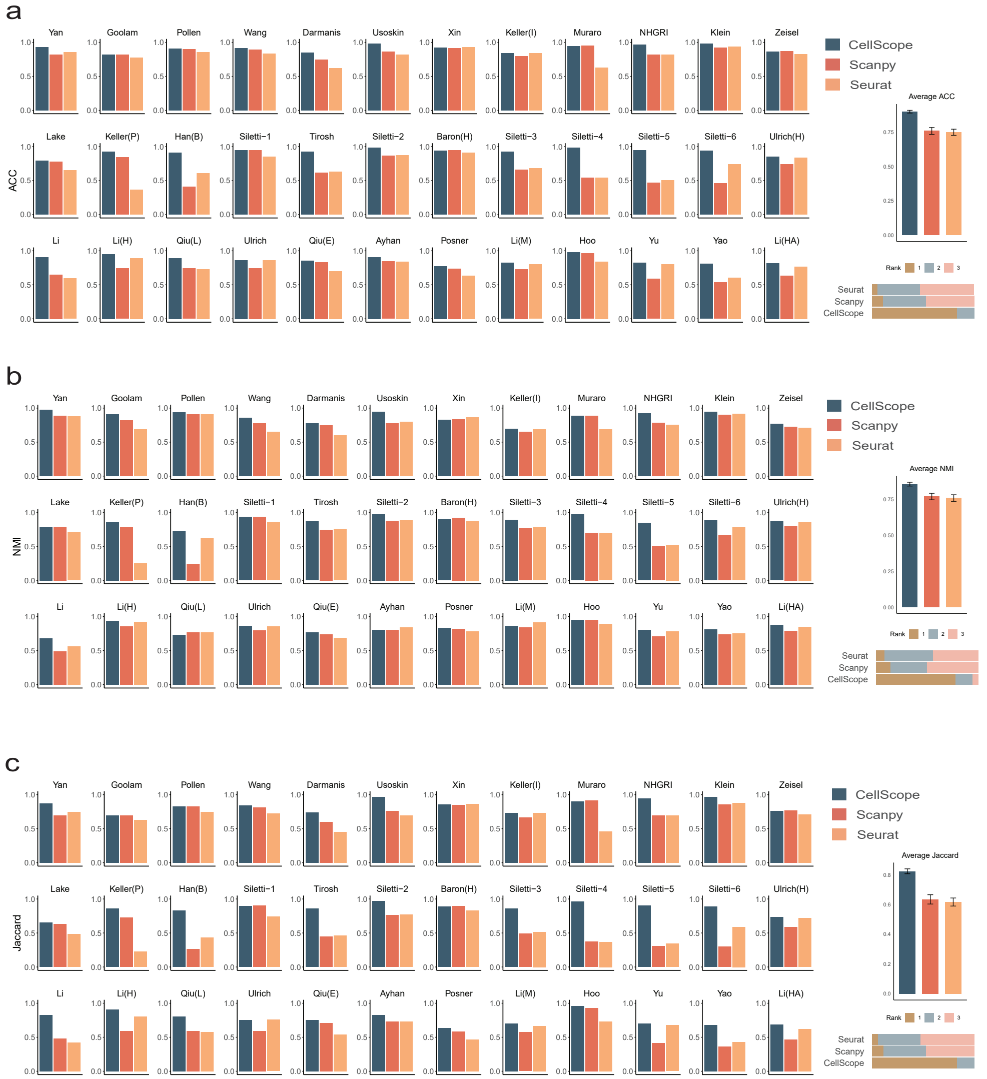

Figure S1: Clustering Performance of CellScope, Seurat, and Scanpy on 36 Benchmark scRNA-seq Datasets. **a.** Clustering performance evaluation of CellScope, Seurat, and Scanpy using the ACC. The first 36 panels display ACC values for individual datasets, while the final panel shows the mean ACC with standard deviation indicated by vertical bars. Bottom shows the Rank distribution of CellScope, Seurat and Scanpy based on clustering performance. **b.** Clustering performance evaluation of CellScope, Seurat, and Scanpy using the NMI. The first 36 panels display NMI values for individual datasets, while the final panel shows the mean NMI with standard deviation indicated by vertical bars. Bottom shows the Rank distribution of CellScope, Seurat and Scanpy based on clustering performance. **c.** Clustering performance evaluation of CellScope, Seurat, and Scanpy using the Jaccard. The first 36 panels display Jaccard values for individual datasets, while the final panel shows the mean Jaccard with standard deviation indicated by vertical bars. Bottom shows the Rank distribution of CellScope, Seurat and Scanpy based on clustering performance.

#### 3.2 Method Integration Analysis

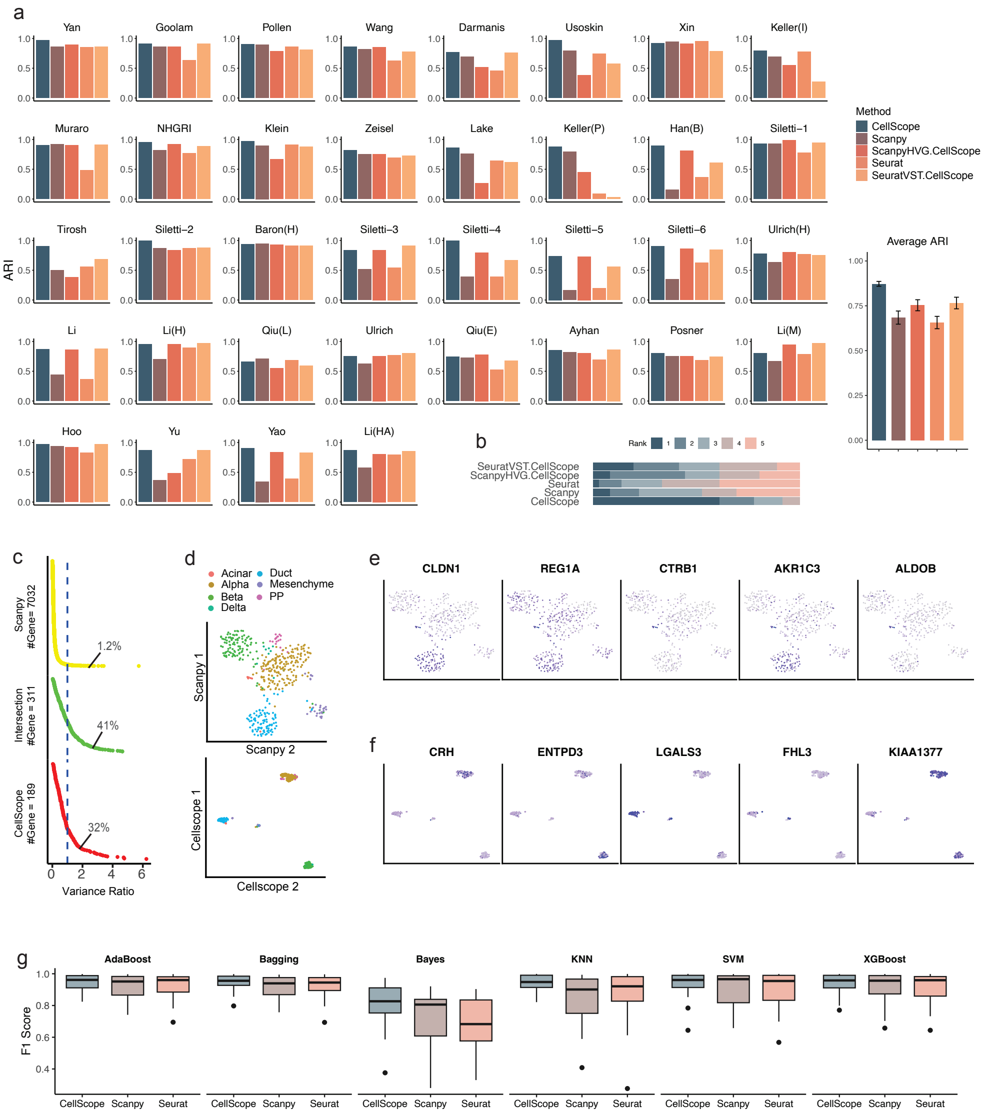

Figure S2: Integration Analysis of CellScope Components with Other Methods. **a.** Clustering performance evaluation of CellScope, Seurat, and Scanpy, CellScope's clustering method with Seurat gene selection(SeuratVST.CellScope) and CellScope's clustering method with Scanpy gene selection(ScanpyHVG.CellScope) using the ARI. The first 36 panels display ARI values for individual datasets, while the final panel shows the mean ARI with standard deviation indicated by vertical bars. **b.** The Rank distribution of CellScope, Seurat, Scanpy, SeuratVST.CellScope and ScanpyHVG.CellScope based on clustering performance. **c.** Gene Selection performance Comparison of CellScope and Scanpy valued by Variance Ratio of Wang Dataset. CellScope can not only capture the high-quality marker genes identified by Seurat but also uncovers meaningful highly variable genes that Scanpy overlooks, the dash line represents variance ratio equals 1. **d.** Venn diagram comparison of gene selection results between CellScope, Scanpy, and Seurat. **e.** Left: Clustering performance of Scanpy on the Usoskin dataset, labeled by cell types and sub-cell types. Right: Clustering performance of Seurat on the Usoskin dataset, labeled by cell types and sub-cell types. **f**. The performance of six supervised classification algorithms on data processed through the gene selection modules of CellScope, Scanpy, and Seurat. The results consistently show CellScope's ability to enhance classification accuracy across various algorithms and datasets

##### 3.3 Gene Expression Analysis: Keller Dataset

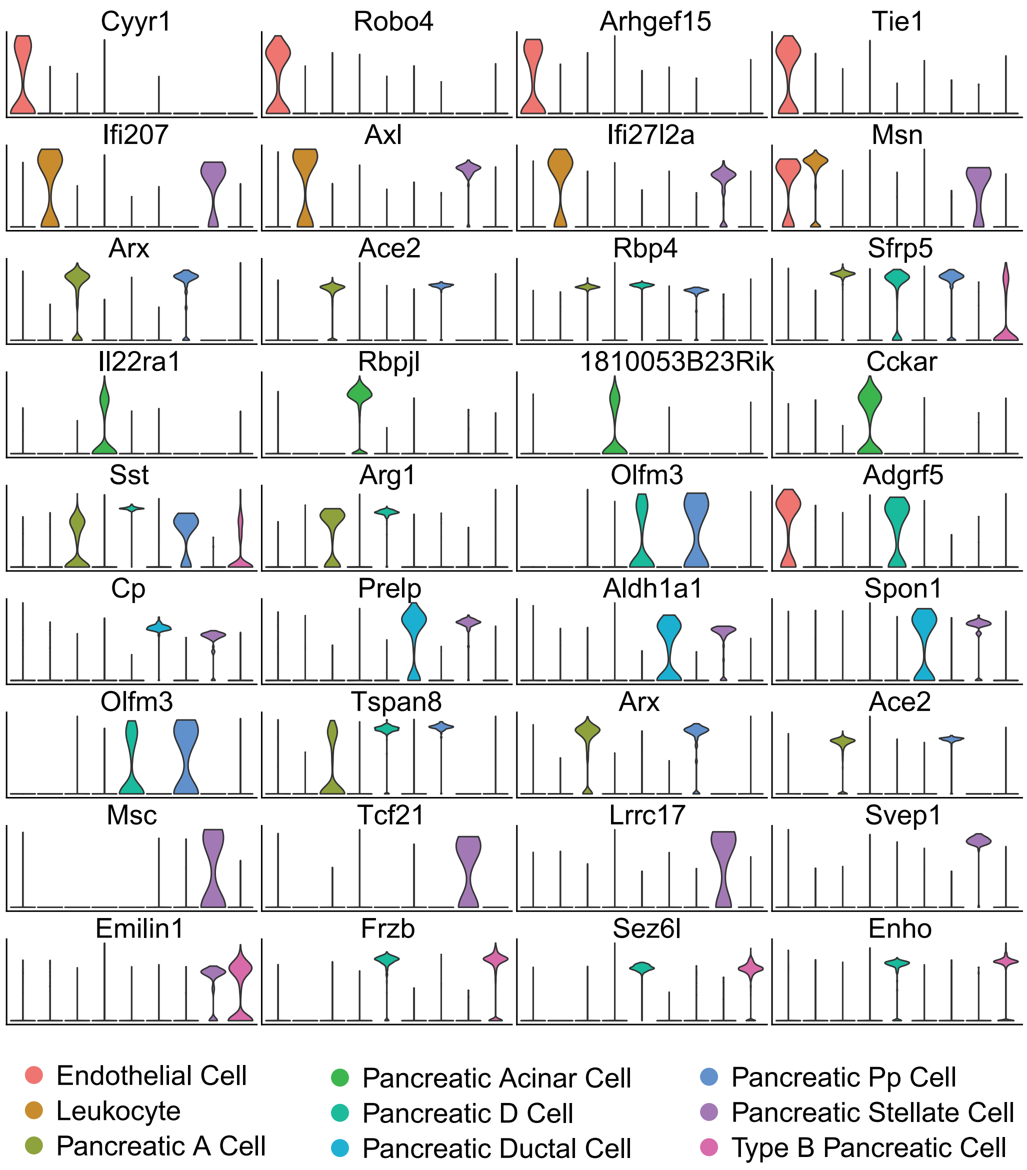

Figure S3: Cell-type Specific Gene Expression Patterns in Keller Dataset Using CellScope. Expression distribution of four marker genes per cell type, identified through CellScope's gene selection method. Each row represents cell-type specific expression patterns, demonstrating the method's capability in identifying distinctive markers for downstream analysis.

#### 3.4 Detailed Analysis of Usoskin Dataset

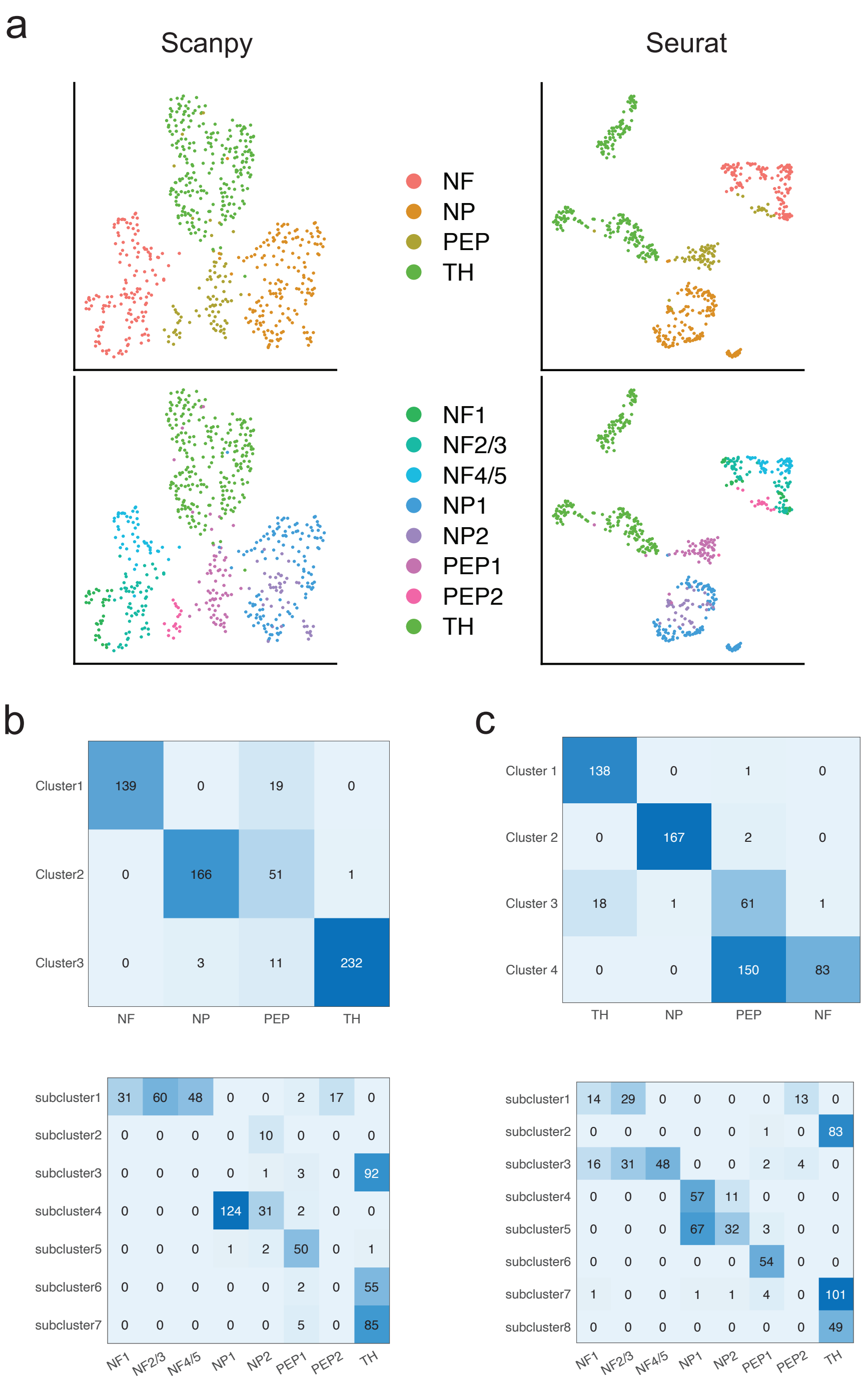

Figure S4: Analysis of Usoskin Dataset Using Scanpy and Seurat. **a.** Visualization comparison between Scanpy (left) and Seurat (right), showing cell types (top) and sub-cell types (bottom). **b.** Confusion matrices of Scanpy clustering results, comparing predicted clusters with true cell types (top) and sub-cell types (bottom). **c.** Confusion matrices of Seurat clustering results, comparing predicted clusters with true cell types (top) and sub-cell types (bottom)

##### 3.5 CellScope Disease-Control Analysis Pipeline

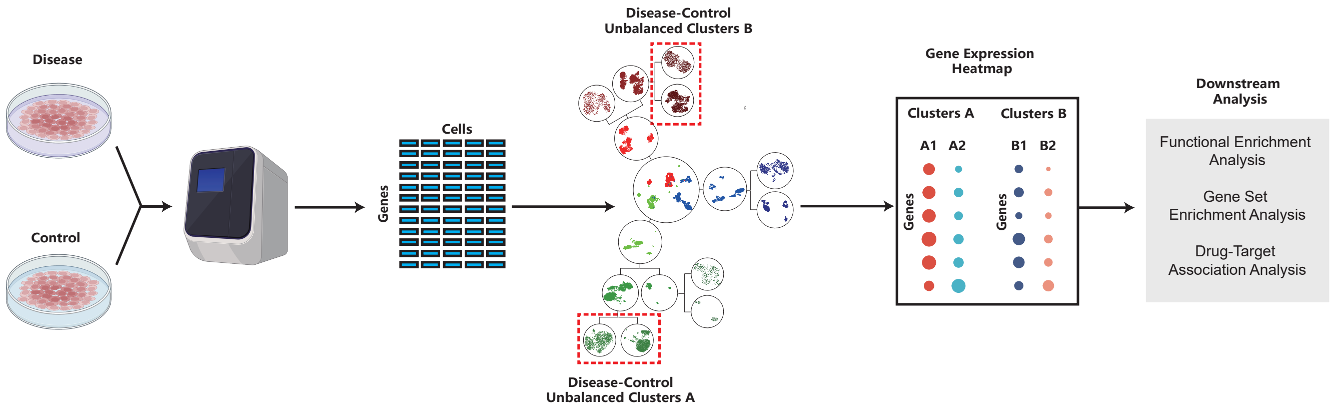

Figure S5: The overview of CellScope-based pipeline for the analysis of disease-control atlas

##### 3.6 Gene Expression Distribution Analysis: Siletti-1 Dataset

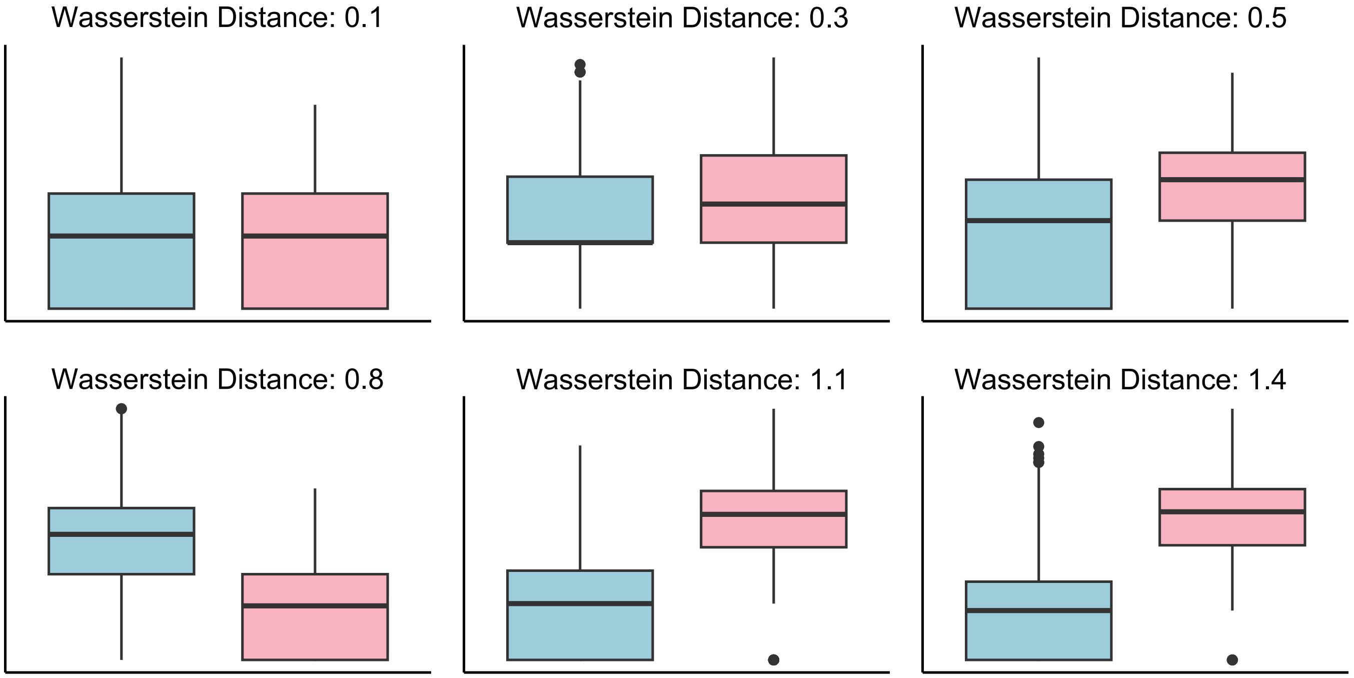

Figure S6: In the Siletti-1 dataset, the gene expression comparison between SubCluster1 and SubCluster2 shows that when the Wasserstein distance is less than 0.5, the gene expression distributions are similar; when the distance is between 0.5 and 1, there is a significant difference in the means, but some overlap remains, indicating moderate differences in gene expression; when the distance exceeds 1, the first quartile of the gene expression boxplot with a higher mean surpasses the third quartile of the other, reflecting substantial differences in gene expression.
